# Evolution of *Wolbachia* Mutualism and Reproductive Parasitism: Insight from Two Novel Strains that Co-infect Cat Fleas

**DOI:** 10.1101/2020.06.01.128066

**Authors:** Timothy P. Driscoll, Victoria I. Verhoeve, Cassia Brockway, Darin L. Shrewsberry, Mariah L. Plumer, Spiridon E. Sevdalis, John F. Beckmann, Laura M. Krueger Prelesnik, Kevin R. Macaluso, Abdu F. Azad, Joseph J. Gillespie

**Affiliations:** Department of Biology, West Virginia University, Morgantown, West Virginia.; Department of Microbiology and Immunology, University of Maryland School of Medicine, Baltimore, Maryland.; Department of Entomology and Plant Pathology, Auburn University, Auburn, Alabama.; Orange County Mosquito and Vector Control District, Garden Grove, California.; Department of Microbiology and Immunology, University of South Alabama, Mobile, Alabama.

**Keywords:** *Wolbachia*, *Ctenocephalides felis*, cat flea, reproductive parasitism, mutualism, lateral gene transfer, cytoplasmic incompatibility, biotin operon

## Abstract

Wolbachiae are obligate intracellular bacteria that infect arthropods and certain nematodes. Usually maternally inherited, they may provision nutrients to (mutualism) or alter sexual biology of (reproductive parasitism) their invertebrate hosts. We report the assembly of closed genomes for two novel wolbachiae, *w*CfeT and *w*CfeJ, found co-infecting cat fleas (*Ctenocephalides felis*) of the Elward Laboratory colony (Soquel, CA). *w*CfeT is basal to nearly all described *Wolbachia* supergroups, while *w*CfeJ is related to supergroups C, D and F. Both genomes contain laterally transferred genes that inform on the evolution of *Wolbachia* host associations. *w*CfeT carries the Biotin synthesis Operon of Obligate intracellular Microbes (BOOM); our analyses reveal five independent acquisitions of BOOM across the *Wolbachia* tree, indicating parallel evolution towards mutualism. Alternately, *w*CfeJ harbors a toxin-antidote operon analogous to the *w*Pip *cinAB* operon recently characterized as an inducer of cytoplasmic incompatibility (CI) in flies. *w*CfeJ *cinB* and immediate-5’ end genes are syntenic to large modular toxins encoded in CI-like operons of certain *Wolbachia* strains and *Rickettsia* species, signifying that CI toxins streamline by fission of larger toxins. Remarkably, the *C*. *felis* genome itself contains two CI-like antidote genes, divergent from wCfeJ *cinA*, revealing episodic reproductive parasitism in cat fleas and evidencing mobility of CI loci independent of WO-phage. Additional screening revealed predominant co-infection (*w*CfeT/*w*CfeJ) amongst *C*. *felis* colonies, though occasionally *w*CfeJ singly infects fleas in wild populations. Collectively, genomes of *w*CfeT, *w*CfeJ, and their cat flea host supply instances of lateral gene transfers that could drive transitions between parasitism and mutualism.

**Importance:** Many arthropod and certain nematode species are infected with wolbachiae which are intracellular bacteria well known for reproductive parasitism (RP). Like other RP strategies, *Wolbachia*-induced cytoplasmic incompatibility, CI, increases prevalence and frequency in host populations. Mutualism is another strategy employed by wolbachiae to maintain host infection, with some strains synthesizing and supplementing certain B vitamins (particularly biotin) to invertebrate hosts. Curiously, we discovered two novel *Wolbachia* strains that co-infect cat fleas (*Ctenocephalides felis*): *w*CfeT carries biotin synthesis genes, while *w*CfeJ carries a CI-inducing toxin-antidote operon. Our analyses of these genes highlight their mobility across the *Wolbachia* phylogeny and source to other intracellular bacteria. Remarkably, the *C*. *felis* genome also carries two CI-like antidote genes divergent from the *w*CfeJ antidote gene, indicating episodic RP in cat fleas. Collectively, *w*CfeT and *w*CfeJ inform on the rampant dissemination of diverse factors that mediate *Wolbachia* strategies for persisting in invertebrate host populations.

## Introduction

Wolbachiae (*Alphaproteobacteria*: Rickettsiales: Anaplasmataceae) comprise Gram-negative, obligate intracellular bacteria that infect over half the world’s described insect species as well as certain parasitic nematodes (1). Unlike other notable rickettsial genera that contain human pathogens (e.g., *Rickettsia*, *Orientia*, *Neorickettsia*, *Anaplasma*, and *Ehrlichia*), wolbachiae do not infect vertebrates (2). A single species, *Wolbachia pipientis*, is formally recognized with numerous members designated as strains within 15 reported supergroups (3–9). Genomic divergence indicates further species names are warranted (10), though increasing diversity and community consensus suggest caution regarding further *Wolbachia* classification at the species level (11, 12).

Like other obligate intracellular microbes, wolbachiae are metabolic parasites that complement a generally reduced metabolism with pilfering of host metabolites (13, 14)). Their ability to survive and flourish is also heavily influenced by the acquisition of key functions through **lateral gene transfer** (**LGT**). Several described *Wolbachia* strains demonstrate characteristics of limited mutualism with their invertebrate partner (15, 16), through the synthesis and provisioning of riboflavin (17) and biotin (18, 19). While riboflavin biosynthesis genes are highly conserved in wolbachiae (20), biotin biosynthesis genes are rare and likely originated via LGT with taxonomically divergent intracellular bacteria (21). Still other strains of *Wolbachia* exert varying degrees of **reproductive parasitism** (RP) on their insect host (22), influencing host sexual reproduction via processes such as male-killing, feminization, parthenogenesis and **cytoplasmic incompatibility** (**CI**) (23). *Wolbachia* genes underpinning CI and male killing have been characterized (24–29) and occur predominantly in the **eukaryotic association module** (**EAM**) of *Wolbachia* prophage genomes (30). These genes highlight the role of LGT in providing wolbachiae with factors facilitating mutualism or RP, both of which are highly successful strategies for increasing infection frequency in invertebrate host populations.

Compared to reproductive parasites, *Wolbachia* mutualists appear to form more stable, long-term relationships with their hosts, as supported by *Wolbachia*-host codivergence in certain filarial nematodes (31) and *Nomada* bees (32). In contrast to the stability of mutualists, relationships of reproductive parasites appear more ephemeral. RP can be a strong mechanism to increase infection frequency, and CI inducing strains can replace populations without infections (33, 34). Despite this, CI is prone to neutralization through the evolution of host suppression (23, 35, 36) and purifying selection on the host doesn’t preserve CI (36). A weakened CI background might be a ripe setting for invasions to begin. Whether invasion occurs at the level of alternate wolbachiae, WO-phages, or CI operons themselves is an area of active evolutionary research, but a clear result of this evolutionary complexity is that RP inducing *Wolbachia* phylogenies are discordant with those of their hosts (37–39). Horizontal transmissions, which can occur through direct host interactions, environmental contacts (shared habitat, food sources, etc.) or predator/parasitoid delivery, contribute to the discordance (40) and possibly drive episodic invasions and replacements.

Horizontal transmission is also evident from hosts infected with multiple *Wolbachia* strains (41–43), though little is known about the dynamics and stability of *Wolbachia* co-infections (44). For instance, in host populations showing variability in single-versus double-infections, does co-infection reflect a transitory phase (i.e. one strain replacing another) or does it reflect stability of both strains in a population? A single infection may be a consequence of closely related strains carrying different arsenals of RP-inducing genes, and hence battling for control of host reproduction (38). A stable co-infection may reflect more divergent strains that utilize different strategies (i.e. nutrient provisioning versus CI) to inhabit different micro-niches in the same host. Such a case is in part witnessed in doubly-infected cochineals (*Dactylopius coccus*), wherein a Supergroup B strain (*w*DacB) carries an arsenal of RP-inducing genes but a Supergroup A strain (*w*DacA) carries none (41). Unfortunately, few systems with divergent co-infecting *Wolbachia* have been characterized sufficiently to dissect contrasting mechanisms that enable long-term host co-infections.

We recently sequenced the genome of the cat flea (*Ctenocephalides felis*) and concurrently closed the genomes of two novel wolbachiae, *w*CfeT and *w*CfeJ, that co-infect fleas of the Elward Laboratory (Soquel, California; hereafter EL fleas) (45). Genome-based phylogeny estimation placed *w*CfeT on an ancestral divergent branch, while *w*CfeJ was found to subtend *Wolbachia* Supergroup C. Both genomes show evidence of past WO phage infection, with LGT hotspots indicating recent acquisition of factors predicted to either underpin mutualism (*w*CfeT) or mediate CI (*w*CfeJ). Additionally, we describe two loci within the *C. felis* genome itself that support repeated acquisitions of CI anti-toxin genes by the host. Infection surveys of cat flea populations and colonies from across the US indicate extensive spread of *w*CfeT among wild populations, and a strong bias for *w*CfeT/*w*CfeJ co-infection in colonies, highlighting the complex ecological relationship that drives this system. Finally, we apply an informatics approach including phylogenomics, synteny analysis, and functional alignment to construct a comprehensive picture revealing the recurrent evolution of nutritional symbiosis and RP across the Rickettsiales.

## Results and Discussion

### Identification of novel *Wolbachia* parasites

Recently, we assembled and annotated closed genomes from two divergent *Wolbachia* strains, named *w*CfeT and *w*CfeJ, that were captured during genome sequencing of the cat flea (EL Colony) (45). The 16S rDNA sequences of these two strains are identical to those previously identified in another cat flea colony (Louisiana State University) (46–48), which historically has been replenished with EL colony fleas. *w*CfeT and *w*CfeJ are broadly similar to other closed *Wolbachia* genomes in terms of length, GC content, number of protein coding genes, and coding density (**Table 1**), although it is interesting to note that *w*CfeT exhibits one of the highest pseudogene counts (second only to *w*Cle) while *w*CfeJ boasts one of the lowest (second only to *w*Ov). Coupled with other recently sequenced genomes for non-Supergroup A and B strains (e.g., *w*Fol and *w*VulC), these cat flea-associated wolbachiae blur the lines demarcating genomic traits previously used to distinguish ancestral *Wolbachia* lineages from those of Supergroups A and B strains (44).

**Table 1.** Summary statistics of selected *Wolbachia* genomes.

| *Taxon* | *RefSeq ID* | *SG* | *Length (nt)* | *GC (%)* | *Protein coding genes* | *Pseudo-genes* | *Mean gene length (nt)* | *Coding density (coding/total nt)* |
| --- | --- | --- | --- | --- | --- | --- | --- | --- |
| wFol | NZ_CP015510.2 | E | 1801626 | 34.4 | 1537 | 85 | 976 | 0.880 |
| wCfeT | NZ_CP051156.1 | ? | 1495538 | 35.2 | 1424 | 201 | 890 | 0.727 |
| wOv | NZ_HG810405.1 | C | 960618 | 32.1 | 650 | 45 | 967 | 0.700 |
| wOo | NC_018267.1 | C | 957990 | 32.1 | 639 | 73 | 968 | 0.719 |
| wCfeJ | NZ_CP051157.1 | ? | 1201647 | 35.6 | 1116 | 51 | 917 | 0.813 |
| wCle | NZ_AP013028.1 | F | 1250060 | 36.3 | 1012 | 210 | 818 | 0.829 |
| wBm | NZ_CP034333.1 | D | 1080064 | 34.2 | 803 | 178 | 868 | 0.789 |
| wBm | NC_006833.1 | D | 1080084 | 34.2 | 839 | 167 | 854 | 0.795 |
| wMau | NZ_CP034334.1 | B | 1273527 | 34.0 | 1045 | 136 | 944 | 0.876 |
| wMau | NZ_CP034335.1 | B | 1273530 | 34.0 | 1047 | 131 | 946 | 0.876 |
| wNo | NC_021084.1 | B | 1301823 | 34.0 | 1066 | 133 | 952 | 0.878 |
| wAlbB FL2016 | NZ_CP041923.1 | B | 1482279 | 34.5 | 1190 | 195 | 917 | 0.858 |
| wAlbB HN2016 | NZ_CP041924.1 | B | 1483853 | 34.4 | 1200 | 187 | 917 | 0.858 |
| wAlbB | NZ_CP031221.1 | B | 1484007 | 34.4 | 1197 | 191 | 916 | 0.858 |
| wMeg | NZ_CP021120.1 | B | 1376868 | 34.0 | 1136 | 125 | 948 | 0.869 |
| wHa | NC_021089.1 | A | 1295804 | 35.1 | 1123 | 104 | 899 | 0.852 |
| wCauA | NZ_CP041215.1 | A | 1449344 | 35.0 | 1260 | 134 | 861 | 0.860 |
| wAna | NZ_CP042904.1 | A | 1401460 | 35.2 | 1218 | 107 | 893 | 0.857 |
| wRi | NC_012416.1 | A | 1445873 | 35.2 | 1245 | 122 | 892 | 0.859 |
| wAu | NZ_LK055284.1 | A | 1268461 | 35.2 | 1099 | 125 | 878 | 0.855 |
| wMel | NC_002978.6 | A | 1267782 | 35.2 | 1116 | 129 | 863 | 0.857 |
| wMel I23 | NZ_CP042444.1 | A | 1269137 | 35.2 | 1119 | 128 | 862 | 0.857 |
| wMel N25 | NZ_CP042446.1 | A | 1267781 | 35.2 | 1120 | 125 | 863 | 0.857 |
| wMel ZH26 | NZ_CP042445.1 | A | 1267436 | 35.2 | 1117 | 125 | 865 | 0.857 |
| wIrr | NZ_CP037426.1 | A | 1352354 | 35.3 | 1232 | 115 | 827 | 0.855 |

Prior reports have indicated the presence of multiple *Wolbachia* “types” infecting *C*. *felis*, with phylogenies inferred from partial gene sequences placing these unnamed *Wolbachia* in supergroups B or F, or in ancestral lineages (49–51). Our robust genome-based phylogeny estimation reveals that both *w*CfeT and *w*CfeJ are early branching wolbachiae (**Fig. 1**; **Table S1**). *w*CfeT is similar to undescribed *C*. *felis*-associated strains that branch ancestrally to most other *Wolbachia* lineages (51–53), while *w*CfeJ is similar to undescribed *C*. *felis*-associated strains closely related to *Wolbachia* supergroups C, D and F (49, 51). The substantial divergence of *w*CfeT and *w*CfeJ from a *Wolbachia* supergroup B strain associated with *C*. *felis* from Germany (*w*Cte) indicates a diversity of wolbachiae that are capable of infecting cat fleas. Importantly, the nature of host-microbe relationships for any of these *C*. *felis*-associated wolbachiae is unknown.

**Figure 1.**
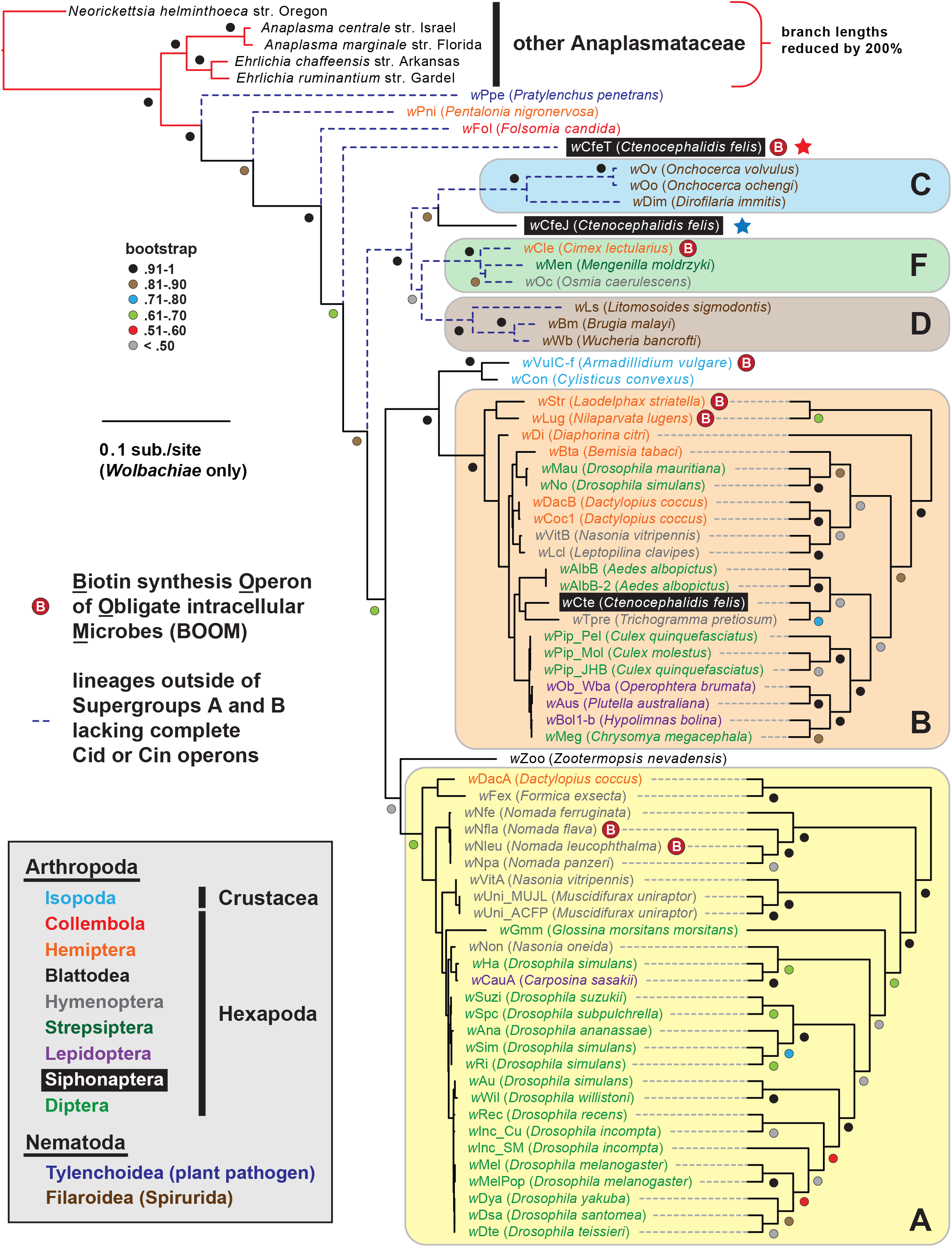
*Wolbachia* genome-based phylogeny estimation. *Wolbachia* supergroups are within colored ellipses. *Ctenocephalides felis*-associated Wolbachiae are within black boxes, with red (*w*CfeT) and blue (*w*CfeJ) stars depicting the two novel *Wolbachia* parasites infecting *C*. *felis*. Gray inset: *w*CfeT and *w*CfeJ genome sequences and assembly statistics. White inset: color scheme for nematode and arthropod hosts. Phylogeny was estimated for 66 *Wolbachia* and five outgroup genomes (see Materials and Methods for more details). Branch support was assessed with 1,000 pseudoreplications. Final ML optimization likelihood was -52350.085098.

### *w*CfeT is equipped with biotin synthesis genes

The enzymatic steps for bacterial biosynthesis of biotin from malonyl-CoA are largely conserved (54) and usually involve six *bio* enzymes and the fatty acid biosynthesis machinery (55) (**Fig. 2A**). We previously identified the first plasmid-borne *bio* gene operon, which occurs on plasmid pREIS2 of *Rickettsia buchneri*, an endosymbiont of the black-legged tick (21). We observed a rare gene order for the pREIS2 *bio* operon shared only with *bio* operons encoded on the chromosomes of obligate intracellular pathogens *Neorickettsia* spp. and *Lawsonia intracellularis* (*Deltaproteobacteria*). These unique *bio* operons formed a clade in estimated phylogenies, indicating LGT of the operon across divergent intracellular species. Subsequent studies, using our same dataset for phylogeny estimation, discovered this *bio* operon in certain *Wolbachia* (19, 32, 56), *Cardinium* (57, 58) and *Legionella* (59) species. We refer hereafter to this intriguing LGT of *bio* genes as the <u>B</u>iotin synthesis <u>O</u>peron of <u>O</u>bligate intracellular <u>M</u>icrobes (BOOM).

**FIG 2.**
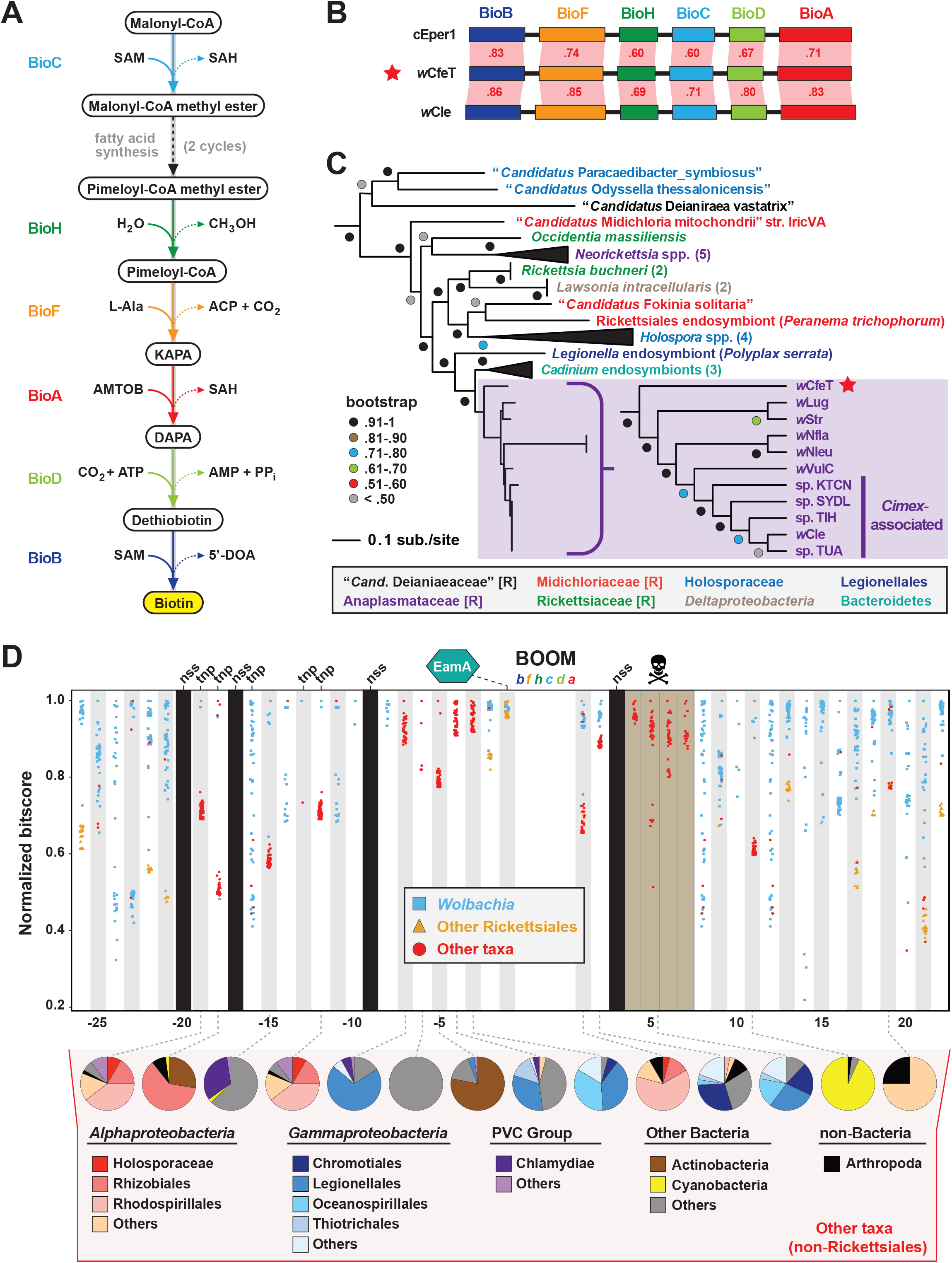
*w*CfeT carries the Biotin synthesis Operon of Obligate intracellular Microbes (BOOM). (A) Metabolism of biotin from malonyl-CoA (55). (B) Comparisons of *w*CfeT BOOM to equivalent loci in *Cardinium* endosymbiont of *Encarsia pergandiella* (cEper1, CCM10336-CCM10341) and *Wolbachia* endosymbiont of *Cimex lectularius* (*w*Cle, BAP00143-BAP00148). Red shading and numbers indicate % identity across pairwise protein alignments (Blastp). Gene colors correspond to enzymatic steps depicted in panel **A**, with all proteins drawn to scale (as a reference, *w*CfeT BioB is 316 aa). (C) Phylogeny of BOOM and other *bio* gene sets from diverse bacteria estimated from the concatenation of six *bio* enzymes (BioC, BioH, BioF, BioA, BioD, and BioB); see **Table S2** for all sequence information, and **Figure S1A** for complete tree estimated under Maximum Likelihood. In taxon color scheme, [R] denotes Rickettsiales, with Holosporaceae as a revised family of Rhodospirillales (101). (D) Analysis of genes flanking *w*CfeT BOOM. See **Table S3** for comparison with other BOOM-containing wolbachiae. Graph depicts top 50 Blastp subjects (e-value < 1, BLOSUM45 matrix) using the *w*CfeT proteins as queries against the NCBI nr database (January 9, 2020). Colors distinguish three taxonomic groups (see inset). Black columns: no significant similarity (nss) in the NCBI nr database. Tnp, predicted transposase. Brown, pseudogenization. See **Figure S1B** for EamA phylogeny estimation. Pie charts across the bottom delineate taxonomic diversity of hits to “Other taxa,” for *w*CfeT proteins with significant Blastp matches outside of the Rickettsiales.

*w*CfeT carries a complete BOOM with greater similarity to *Wolbachia* BOOM than those of other bacteria (**Fig. 2B**). Phylogeny estimation of BOOM and other diverse *bio* gene sets indicates *w*CfeT BOOM is ancestral to other *Wolbachia* BOOM (**Fig. 2C, Fig. S1**). However, as the divergence pattern for other *Wolbachia* BOOM is discordant with genome-based phylogeny (**Fig. 1**), it is clear that multiple independent BOOM acquisitions have occurred in wolbachiae irrespective of lineage divergences. This is supported by a previous study hypothesizing recent BOOM acquisition in *w*Nfla and *w*Nleu (32), as well as the nature of BOOM flanking regions in the genomes of *w*CfeT, *w*VulC, *w*Cle, *w*Str, *w*Lug, which collectively lack synteny and are riddled with transposases, recombinases and phage-related elements (**Table S3**). Furthermore, analysis of *w*CfeT BOOM flanking genes reveals hotspots for LGT, exemplified by an S-adenosylmethionine importer (EamA) gene (**Fig. S1B**) that is a signature among many diverse obligate intracellular bacteria (21, 60–62).

As some BOOM-containing *Wolbachia* strains have been shown to provide biotin to their insect hosts (18, 19), we posit that *w*CfeT has established an obligate mutualism with *C*. *felis* mediated by biotin-provisioning. This is consistent with a WO prophage in the *w*CfeT genome that is highly colinear to the WOVitA1 prophage (30), except for the absence of a EAM and hence no RP-inducing loci (**Fig. S2**). *Wolbachia* strains associated with bedbug (*w*Cle), planthopper (*w*Str, *w*Lug), and termites (*w*Ctub) with evidence for BOOM (63) reside in bacteriocytes of their insect hosts, structures associated with nutritional symbiosis in many arthropods (64). Interestingly, *C*. *felis* is not known to contain bacteriocytes, and the ability of *w*CfeT to synthesize and supplement biotin to cat fleas awaits experimentation.

### *w*CfeJ lends insight on the evolution of reproductive parasitism

*w*CfeJ carries a **toxin-antidote** (**TA**) operon that is architecturally similar to the *w*Pip_Pel CinA/B TA operon (**Fig. 3A**), which was previously characterized as a CI inducer in flies (24). Grouped into three types (26, 65), CinB toxins harbor dual **nuclease** (**NUC**) domains in place of the **deubiquitinase** (**DUB**) domain of CidB, another CI inducer in flies (26–28). Chimeric NUC-DUB toxins, termed CndB toxins (66), also occur as operons (*cndAB*) in certain *Wolbachia* and *Rickettsia* genomes (27, 67). Although their functions remain unknown, these larger CndB toxins belong to an extraordinarily diverse array of toxins with highly modular architectures that occur in divergent intracellular bacteria (67). Curiously, despite a size similar to the CinB toxins, *w*CfeJ’s CinB toxin has much higher sequence identity to larger CndB and NUC-containing toxins found mostly in *Wolbachia* Supergroups A and B (**Fig. 3B**). A similar pattern was found for *w*CfeJ’s CinA antidote, indicating the genes were likely acquired as an operon.

**FIG 3.**
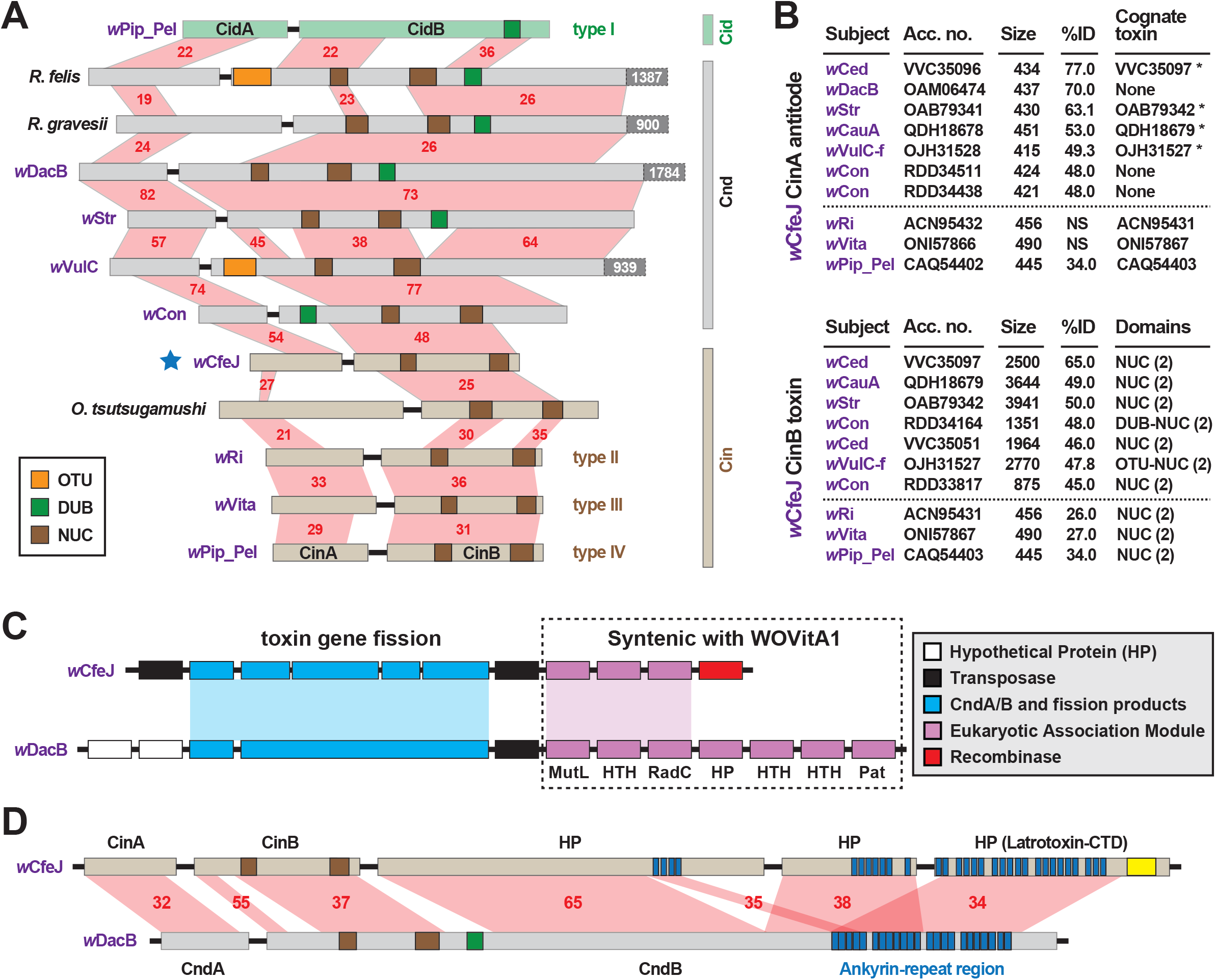
*w*CfeJ lends insight on the evolution of reproductive parasitism. (A) Comparison of known and predicted RP-inducing toxin-antidote operons of diverse rickettsial species. Light green: CidA/B of wPip_Pel (CAQ54390, CAQ54391); grey: CndA/B of pLbAR_36/38 (KHO02168, KHO02170), *Rickettsia gravesii* (WP_017442886, WP_024547315), *w*DacB (OAM06111, OAM06112), *w*Str (OAB79364, OAB79365), *w*VulC (OJH31528, OJH31527), *w*Con (RDD34163, RDD34164); light brown, CinA/B of *w*CfeJ (blue star, WP_168456002 & WP_168456001), *Orientia tsutsugamushi* (CAM80637, CAM80639), *w*Ri (ACN95432, ACN95431), *w*Vita (ONI57866, ONI57867), and wPip_Pel (CAQ54402, CAQ54403). Type I-IV CI operons of *Wolbachia* parasites (26) are distinguished. Protein domains as follows: green, CE clan protease; brown, PD-(D/E)XK nuclease; orange, OTU cysteine protease. Red shading and numbers indicate % identity across pairwise alignments. Proteins are drawn to scale, with additional C-terminal sequence for some CndB toxins depicted in dark gray boxes. (B) Sequence similarity profiles for *w*CfeJ’s CinA antidote (**top**) and CinA toxin (**bottom**). Subjects are from a Blastp search against the NCBI ‘Wolbachia’ database. Size, amino acids. Antidotes with cognate toxins (operon partners) also recovered in searches are noted with asterisks. Top seven hits are shown, with matches (where significant) to proteins of types II-IV TA operons shown below dashed lines. Toxin domains are from our prior report or predicted newly here using the EMBL’s Simple Modular Architecture Research Tool (SMART) (174). Only NUC and DUB domains are listed, though larger toxins contain many other predicted domains. (C) Comparison of *w*CfeJ and *w*DacB degraded WO prophage genomes captures streamlining of a CI-like TA operon from a Cnd operon. WO prophage coordinates: *w*CfeJ, WP_168455994-WP_168456003; *w*DacB, OAM06111-OAM06118 (LSYY01000098). Schema follows the description of the WOVitA prophage (30). (D) *w*CfeJ’s CinB-like toxin evolved from gene fission of a larger CndB-like toxin. NCBI protein accession numbers: *w*CfeJ (WP_168455999, WP_168456000, and WP_168456001); *w*DacB (OAM06111, OAM06112). Red shading and numbers indicate % identity across pairwise alignments. Proteins are drawn to scale; as a reference, *w*DacB CndB is 3707 aa.

In contrast to *w*CfeT, *w*CfeJ harbors a highly degraded WO prophage genome with only a partial EAM, the CinA/B operon, and several other genes encoding hypothetical proteins and transposases (**Fig. 3C**). This is reminiscent of the *w*Rec parasite, which lacks a functional WO phage yet possesses TA modules within remnant EAMs (68). Synteny analysis with other *Wolbachia* phage revealed *w*CfeJ *cinB* and three downstream genes are colinear with a large (3707 aa) *w*DacB *cndB* toxin (**Fig. 3D**). This provides strong evidence for the fission of a large modular CndB toxin into a smaller CinB toxin. This interpretation is also consistent with our prior hypothesis stating that more streamlined RP-inducing toxins (i.e., *cinB* and *cidB*) originate from larger modular toxins (i.e., *cndB* and others) that are widespread in the intracellular mobilome, particularly *Wolbachia* phage and *Rickettsia* plasmids and **integrative and conjugative elements** (**ICEs**) (48, 67).

*w*CfeJ is unique among described early-branching wolbachiae by containing a CI-like TA operon. In fact, the *Wolbachia* strains in closely related Supergroups C, F and D are considered mutualists with their nematode and arthropod hosts. *w*Fol, another ancestral *Wolbachia* lineage (**Fig. 1**), does carry a myriad of large modular candidate RP-inducing toxins but does not have *cin*, *cid* or *cnd* TA operons (69, 70). Collectively this indicates acquisition of a WO prophage by *w*CfeJ, possibly from another Supergroup A or B *Wolbachia* strain that also infects cat fleas (e.g., *w*Cte), with viral or *Wolbachia* level selection streamlining a *cndAB* locus into a *cinAB* operon. Given that the genomes of many *Wolbachia* reproductive parasites harbor diverse arrays of CinA/B-and CidA/B-like operons (66, 67), we posit that *w*CfeJ’s CinA/B TA operon might function in CI or some other form of RP; however, further study is required to evaluate *w*CfeJ as a cat flea reproductive parasite. Furthermore, an intriguing speculation is that functions of the larger modular toxins, exemplified by *w*DacB CndB, whatever they be, are likely the primal functions that gave rise to CI.

### Lateral transfer of *Wolbachia* CI-like antidotes to the cat flea genome

Our previous work identified CI-like toxin and antidote genes on scaffolds in several arthropod genome assemblies, although antidote genes were found more frequently possibly due to utility in suppressing microbial invaders implementing RP (67). Many of these genomes contain multiple divergent antidote genes, though none were found to harbor large-scale *Wolbachia* gene transfer that is known to occur in certain invertebrates (71–77). In light of this observation, we originally speculated that CI-like genes are mobilized from wolbachiae to host genomes possibly through WO phage introgression; however, the absence of other WO phage genes in these genomes made this assertion tenuous. Instead, more recent work describing the first WO prophage carrying two sets of CI loci provides compelling evidence for transposon-mediated LGT of CI loci between divergent wolbachiae (38). This phage-independent mechanism for RP gene purging and replenishment in wolbachiae may result in the inadvertent capture of these genes by host genomes.

When we analyzed the cat flea genome for CI-like genes, we discovered that the largest *C*. *felis* scaffold (185.5 Mb) was found to harbor two CI-like antidote genes (**Fig. 4A**). Remarkably, neither antidote has any significant similarity to CinA of *w*CfeJ (**Fig. 4B**), indicating that these CI antidotes originated from other, as yet undescribed wolbachiae. The *C. felis* CinA-like CDS (XP_026474240.1) has greater similarity to CinA proteins than CidA proteins, while the CidA-like CDS (not annotated in the current *C*. *felis* assembly) has much greater similarity to CidA antidotes (27, 28, 78–80). The *C*. *felis cinA*-like gene is found as an exon within a larger flea gene model (LOC113377978), and is well supported (18X coverage, 7.2 TPM) by transcripts from the 1KITE project (81). Since these transcripts were generated in fleas from a different colony (Dr. Michael Dryden, Kansas State University), we conclude that this gene transfer occurred during an historical infection predating the establishment of these two colonies. Finally, the *C*. *felis* CinA- and CidA-like antidotes each possess the conserved motifs we previously defined for all CI antidotes (67), with the exception of a truncation of motifs 5 and 6 in the CinA-like protein (**Fig. 4D**). As other arthropod-encoded antidotes share this C-terminal truncation, it may signify an expendable domain in the host (i.e., a bacterial secretion signal, a motif involved in cognate toxin activation or delivery, etc.).

**FIG 4.**
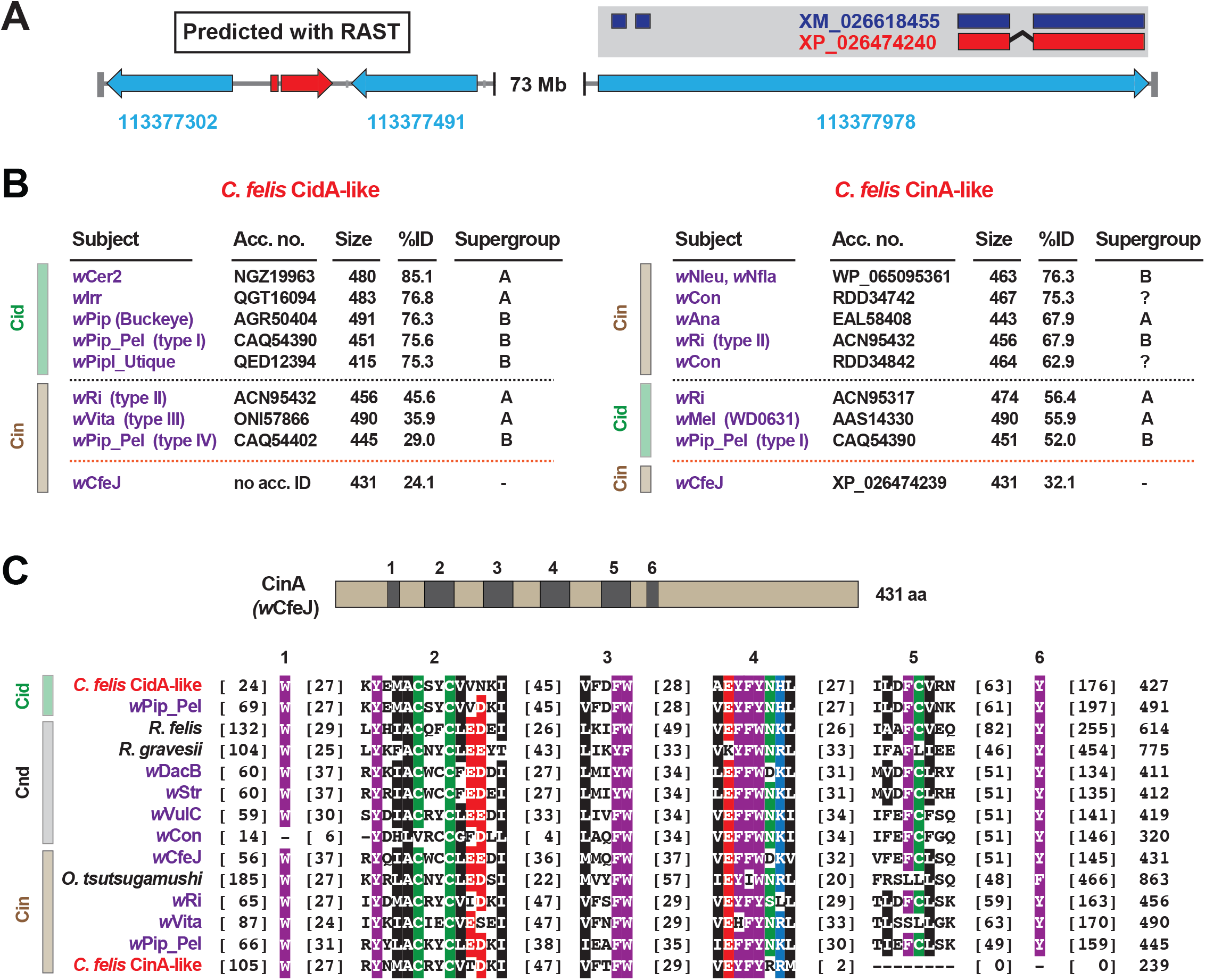
Identification of two CI-like antidote genes in the cat flea genome. (A) The largest *C*. *felis* scaffold (NW_020538040) contains a CidA-like gene, predicted using RAST (185) (**left**), and a CinA-like exon encoded within a larger gene model with transcriptional support (**right**). Red, CI-like antidote genes. Blue, *C*. *felis* genes: 113377302 (RING finger and SPRY domain-containing protein 1-like); 113377491 (negative elongation factor B); 113377978 (uncharacterized). The CinA-like exon (XP_026474240) contains a corresponding transcript (dark blue, XM_026618455) generated from a study (81) independent of the cat flea genome sequencing (45). (B) Sequence similarity profiles for the *C*. *felis* CidA-like (**left**) and CinA-like (**right**) antidotes. Subjects above the black dashed line are the top hits from a Blastp search against the NCBI nr protein database; subjects below are weaker matches to CinA (**left**) and CidA (**right**) proteins; matches to *w*CfeJ CinA are below the orange line. Size, amino acids. (C) Alignment of *C*. *felis* CidA-like and CinA-like antidotes to CidA, CndA and CinA proteins from selected Rickettsiaceae species and *Wolbachia* strains. Schema at top depicts the *w*CfeJ CinA antidote and location of six conserved regions shown in the alignment below. Amino acid coloring as follows: black, hydrophobic; red, negatively charged; green, hydrophilic; purple, aromatic; blue, positively charged. Alignment generated using MUSCLE, default parameters (169).

The incorporation of a *Wolbachia* antidote into an exon of a host gene provides one explanation for how such a laterally transferred gene would remain functional in the absence of the native regulatory elements encoded on WO prophage genomes. We previously hypothesized that CI antidote genes would be under strong selection in arthropod genomes if their expression could result in rescuing CI induction by reproductive parasites (67). In support of this supposition, it has been shown that: host genotypes modulate CI (68, 82–84); some hosts show weak (*Dmel* and *D*. *suzukii* (35, 85, 86)) or strong (*Culex pipiens* and *D*. *simulans* (87, 88)) CI; and a *Wolbachia* strain’s CI can be suppressed when transfected across different hosts (84). While none of the arthropod-encoded CI antidotes have been characterized, we propose they impart immunity to toxins secreted by intracellular parasites, curtailing chronic parasite drive into populations. At least four independent evolutionary mechanisms might lead to suppression of CI. Gene family expansion of, and/or overexpression of key factors capable of counteracting CI mechanisms is possible; similarly, it was postulated that evolution in Cid interacting proteins like karyopherin-α or P32 could lead to target site resistance (25). A third possibility contributing to weakening CI, though not suppression per se, is simply the mutational collapse of the operons themselves due to lack of purifying selection (89). We supply evidence of a fourth possibility whereby CI might be counteracted by germline expression of acquired CI antidotes. Significantly, our finding raises awareness of a potential barrier to *Wolbachia*-mediated pathogen biocontrol measures.

### Characterization of *w*CfeT and *w*CfeJ infection of cat fleas

Coupled with earlier reports indicating multiple early-branching wolbachiae lineages infecting *C*. *felis* (49, 51), our characterization of two divergent *Wolbachia* strains infecting cat fleas in the EL colony (45) as well as fleas maintained for a decade at LSU (46–48) prompted us to test individual fleas for double infection. In addition to EL fleas, three other colony strains and three geographically diverse wild cat flea populations were also surveyed (**Fig. 5A**). Aside from the Modesto Wild strain, which is historically replenished with wild-caught cat fleas (Dr. Bill Donahue, personal communication), strong *w*CfeJ-*w*CfeT co-infection was found in colony fleas. Conversely, single *w*CfeT infection dominated wild cat fleas, with only a few fleas from a large sampling throughout Orange Co. CA harboring *w*CfeJ. While far greater sampling of both colony and wild populations is necessary, our results hint at a strong selection for *w*CfeT over *w*CfeJ in nature. The ability of *w*CfeJ to highly infect colony fleas may arise as a consequence of artificial bottlenecks within colonies, which are known to strengthen CI and maternal transmission (36). Nonetheless, the ability of *w*CfeT and *w*CfeJ to co-infect individual cat fleas provides a system to study contrasting forces (nutritional symbiosis and RP) that potentially drive multiple *Wolbachia* infections in a single invertebrate host.

**FIG 5.**
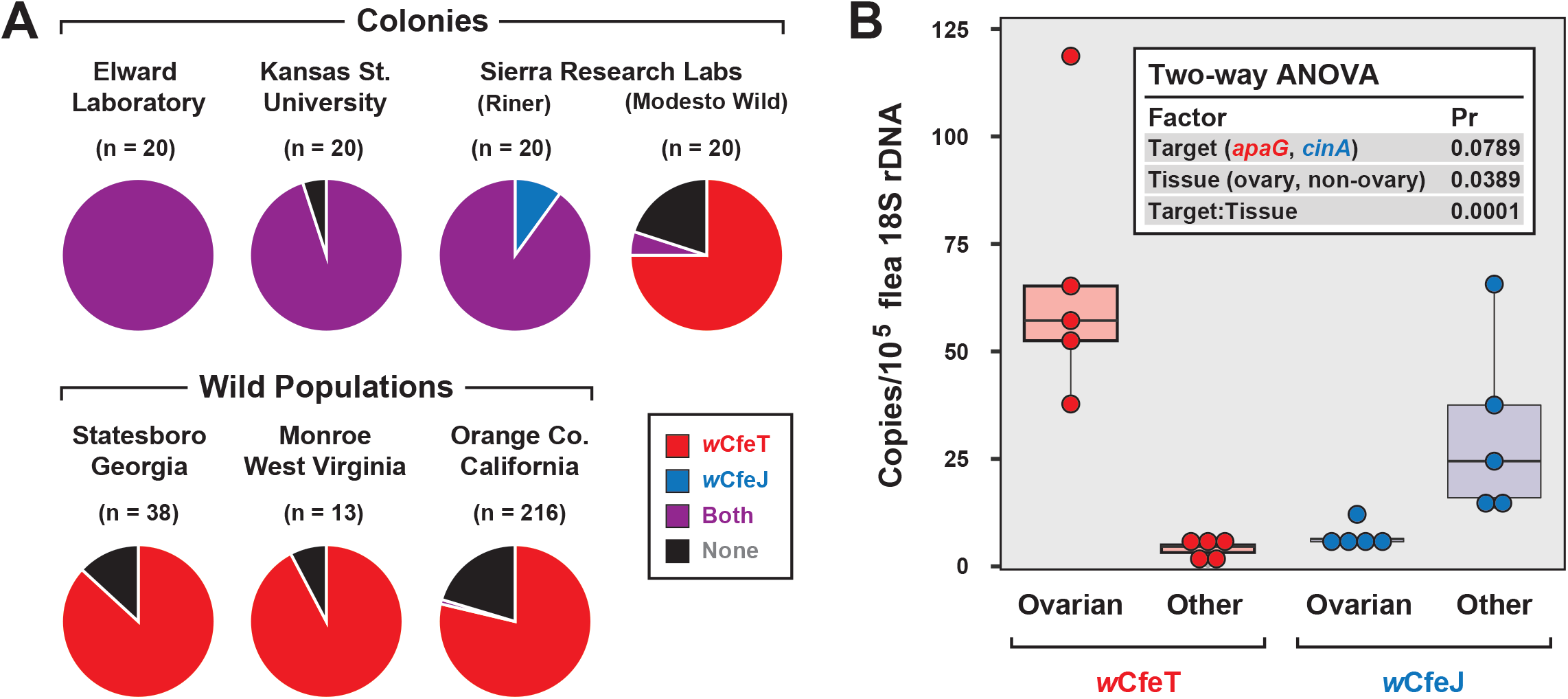
Characterization of *w*CfeJ and *w*CfeJ in cat fleas. (A) Screening for wolbachiae in geographically diverse cat flea colonies and wild populations. Cat fleas from six locations (including two strains from Sierra Research Laboratories) were assessed for the presence of *w*CfeT and *w*CfeJ. *Wolbachia* infection of individual fleas was assessed by qPCR (see **Materials and Methods** for details). (B) Localization of *w*CfeT and *w*CfeJ in cat flea tissues. Dissected ovaries and other tissues (mostly midguts) from individual cat fleas (EL Colony) were assayed by qPCR see **Materials and Methods** for details). Five fleas were assayed for each tissue type.

To gain more insight into the nature of the *w*CfeT-*w*CfeJ co-infection, we analyzed the distribution of *w*CfeT and *w*CfeJ in EL flea tissues (**Fig. 5B**). Surmising that both *Wolbachia* strains are maternally transmitted, we dissected out the ovaries of unfed virgin female fleas and compared *Wolbachia* density in these tissues versus others (primarily midgut). Counter to what we expected, *w*CfeT predominantly localizes in flea ovaries, while the CI harbinger, *w*CfeJ, was predominantly in somatic tissues. Although the two are not mutually exclusive and they do co-localize to some extent. The significance of this distribution is unclear. *w*CfeJ may relocate to ovaries upon a mating cue, or perhaps exert CI via some undefined somatic cell mechanism. Transmission of *w*CfeJ may also be predominantly horizontal via environmental or direct flea contact (per (40)). Sampling co-infected fleas at specific timepoints during development, as well as determining *w*CfeJ tissue localization in the absence of *w*CfeT infection, will be required to unravel the intertwined transmission dynamics of these *Wolbachia* strains in cat fleas.

### Evolution of *Wolbachia* Mutualism and Reproductive Parasitism

Order Rickettsiales is a highly diverse group of obligate intracellular bacteria (2, 62, 90). A rapid pace of newly identified lineages (60, 91–104) provides a rich source for evaluating mechanisms of evolution within constraints of the eukaryotic cell. Phage, plasmids, transposons, ICEs, and other insertion sequences are variably present across rickettsial genomes and mobilize numerous genes to offset reductive genome evolution and provide lifestyle altering traits (21). Our analyses of *Wolbachia* BOOM and RP genes prompted us to evaluate the recurrent evolution of nutritional-mediated mutualism and reproductive parasitism across diverse rickettsial lineages (**Fig. 6**).

**FIG 6.**
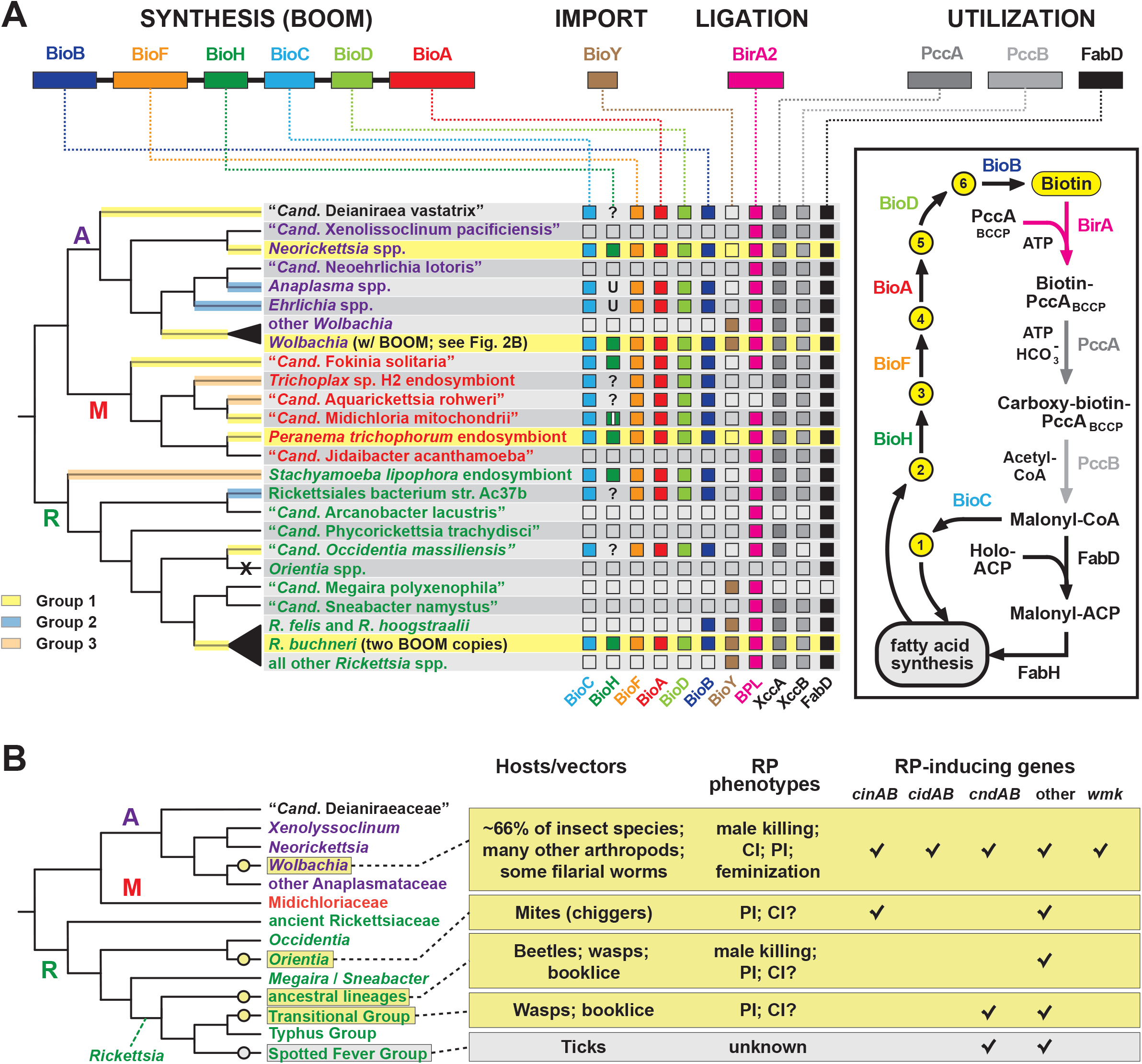
Recurrent evolution of nutritional-mediated mutualism and reproductive parasitism across diverse rickettsial lineages. (A) Diverse strategies for biotin biosynthesis, acquisition and utilization in Rickettsiales. **Top**, biotin synthesis genes (BOOM arrangement), the BioY biotin transporter, the biotin ligase BirA, and biotin utilization enzymes (biotin-dependent propionyl-CoA carboxylase complex and FabD) involved in malonyl-CoA synthesis. The distribution of these genes across major rickettsial lineages is summarized, with phylogeny of Rickettsiales drawn as a consensus from recent studies (13, 91, 101, 103). The acquisition of biotin synthesis operons is depicted by yellow, blue and orange highlighting over branches (see estimated phylogeny in Figure 1A). Taxa highlighted yellow indicate the conservation of BOOM gene arrangement (see **Figure S3**). Question marks indicate missing BioH enzymes in otherwise complete biotin synthesis pathways. These genomes were scanned for other functionally equivalent methyl esterases and found to contain none (**Figure S4D**). Inset highlights the gatekeeper role of BioH in shunting Pimeloyl-CoA methyl ester (metabolite 2) away from fatty acid biosynthesis for the production of biotin. All other metabolites in the biotin synthesis pathway are described in Figure 2A. X, complete loss of biotin utilization in *O*. *tsutsugamushi*. (B) Phylogenomic analysis of RP in Rickettsiales. Phylogeny estimation is further simplified relative to the tree in panel **A**. RP is considered an acquired trait, with origins shown as yellow circles. Spotted Fever Group rickettsiae are shown in gray since RP is unknown, though CI-like genes are present in some species. Information on rickettsial hosts/vectors (2) and known RP (67, 186) are compiled from numerous reports. Check marks reflect the compilations of groups of RP-inducing genes from our prior report, or the recent identification of the *WO-mediated killing* (*wmk*) gene implicated in *Wolbachia* male-killing (29). Helix-turn-helix domains of certain XRE-family like proteins across Rickettsiales were not considered related to *wmk*-encoded proteins. ‘Other’ refers to additional proteins presented in our prior report (67).

#### Nutritional-mediated mutualism

Phylogenetic estimation supports three independent acquisitions of biotin biosynthesis genes by rickettsial species (**Fig. 6A**, **Fig. S1A**). The largest clade (group 1) contains all BOOM-containing species, though only *Neorickettsia* spp., certain wolbachiae, *R*. *buchneri*, and the *Peranema trichophorum* endosymbiont carry the conserved BOOM gene arrangement (highlighted yellow in **Fig. 6A**). Other species in this group, as well as Groups 2 and 3, show different gene arrangements indicating recombination and/or possible replacement of specific *bio* genes through additional LGT (**Fig. S3**). BOOM and other *bio* gene clusters have repeatedly lost *bioH*, which encodes the methyl esterase generating pimeloyl-CoA, the last step in the synthesis of the pimelate moiety of biotin. Bacteria have evolved numerous strategies for catalyzing this reaction (54, 105–112). Interestingly, a novel methyl esterase of *Ehrlichia chaffeensis* (BioU) was recently shown to complement an *E*. *coli ΔbioH* mutant (113); however, we determined that BioU orthologs are present in many rickettsial species, including some that harbor a BioH already and others with no *bio* synthesis genes at all (**Fig. S4A-C**). This indicates that while BioU may generate pimeloyl-CoA in certain rickettsial species – namely, those with complete *bio* synthesis modules except for BioH – it is likely to have a broad range of functions in bacteria (**Fig. S4D**). Furthermore, some species lack the genes for both BioH and BioU, indicating that alternative pathways for generating pimeloyl-CoA await discovery.

Analysis of the genomic distribution of biotin synthesis genes, the biotin transporter BioY, the biotin ligase BirA, and the enzymes involved in malonyl-CoA synthesis (biotin-dependent propionyl-CoA carboxylase complex and FabD) revealed several strategies for biotin utilization in Rickettsiales (**Fig. 6A**). Species that harbor a complete set of *bio* genes (as discussed above) also contain ligase and utilization enzymes, although (not surprisingly) most do not carry the BioY importer. Genera *Wolbachia* and *Rickettsia* largely rely on parasitic scavenging of host biotin to fuel fatty acid biosynthesis: they are dominated by species that lack *bio* genes but carry the BioY importer, BirA ligase, and utilization enzymes. Two species of Transitional Group Rickettsiae (*R*. *felis* and *R*. *hoogstraalii*) may be able to utilize imported dethiobiotin as well, as they retain the biotin synthase BioB; alternatively, this may represent the frayed end of a decaying metabolic pathway similar to the glycerol phospholipid and folate biosynthesis pathways in Rickettsiae (13, 114). While characterized as two-component transporters (with BioM/N), BioY subunits alone can efficiently transport biotin into cells (115) and also participate with host transporters to relocate cytosolic biotin to vacuoles occupied by pathogens (116). However, certain rickettsial species lacking synthesis genes also lack BioY importers yet retain the ligase and utilization enzymes, indicating the presence of alternative transport mechanisms. Remarkably, *Orientia* spp. lack all genes involved in biotin synthesis, import, ligation and utilization, exhibiting the ultimate form of parasitism for host-dependent fatty acid synthesis: these Rickettsiae must pilfer malonyl-CoA from the host (117), using a transport system yet to be described.

The acquisition of BOOM by certain wolbachiae and *R*. *buchneri* provides a theoretical framework for how LGT facilitates a transition from metabolite thievery to nutritional-mediated symbiosis. However, it is important to consider that these endoparasites may be utilizing the BOOM primarily for selfish purposes, as almost all bacteria require biotin. For example, *Neorickettsia* spp. and the deltaproteobacterium *Lawsonia intracellularis* are pathogens that also harbor the BOOM, suggesting a more complex ecological role surrounding biotin acquisition. If *Neorickettsia* spp. do indeed actively provision biotin to their trematode and insect vectors, as has been shown for *w*Cle (19), such host-specific supplementation would necessitate as-yet unidentified genes encoding export factors as well as stringent regulation of the BOOM.

Canonical regulation of biotin synthesis is accomplished via a repressor, fused to the N-terminal domain of the biotin ligase gene, which down-regulates transcription of the *bio* gene cluster in response to high cellular biotin (118, 119). Interestingly, all rickettsial BirA biotin ligases lack this repressor domain, indicating that alternative mechanisms for regulating biotin synthesis and host supplementation are utilized by these bacteria. “*Candidatus* Aquarickettsia rohweri” and the *Trichoplax* sp. H2 endosymbiont (Midichloriaceae) lack BirA ligases altogether (**Fig. 6A**), and no other biotin ligases were found in these genomes (data not shown). Regarding the *Trichoplax* sp. H2 endosymbiont, our prior analysis of the closely related **Rickettsiales endosymbiont of *Trichoplax adhaerens*** (**RETA**) discovered two biotin-protein ligase N-terminal domain-containing proteins (BPL_N) encoded in the *T*. *adhaerens* genome (62). These host-encoded genes contain introns and eukaryotic regulatory elements yet are strongly supported as rickettsial in origin. Homologs of these *T*. *adhaerens* proteins are also present in the *Trichoplax* sp. H2 genome (120) (NCBI acc. nos. RDD42697 and RDD42436), supporting their cross-domain LGT. This raises the possibility for a metabolic interdependence involving biotin synthesis and utilization in placozoans and their rickettsial symbionts, similar to the mosaic metabolic networks underpinning nutritional symbioses in other systems (121–123). For example, the nested tripartite symbiosis between *Planococcus citri* mealybugs, *Tremblaya princeps*, and *Moranella endobia* shows evidence for LGT of several *bio* genes to the mealybug genome (124); these genes share highest similarity to *Wolbachia* BOOM counterparts (data not shown). This is reminiscent of the pea aphid-*Buchnera* symbiosis, wherein several rickettsial LGTs have been implicated in facilitating metabolic interdependence between host and symbiont (125). For rickettsial endosymbionts of placozoans, host acquisition of BPL_N genes (among others (62)) may have established an inextricable obligatory relationship (126, 127) and may hint at the evolutionary fate of other BOOM-containing rickettsiae. However, while our analyses of the BOOM and other *bio* genes in rickettsial genomes are suggestive, in the absence of experimental validation we caution against implicating these genes as factors underpinning nutritional-mediated mutualism.

#### Reproductive parasitism

Among other bacterial reproductive parasites of arthropods (e.g., species of *Cardinium*, *Arsenophonus* and *Spiroplasma*) wolbachiae are perhaps the best-studied for their roles in RP induction (23). Yet within the Rickettsiales, wolbachiae do not walk alone in inducing RP in their hosts. Based on a survey of the literature and analysis of genomic data for the ever-growing pool of RP-inducing genes, we identify at least five strategies for RP induction that have evolved throughout rickettsial evolution, with four independent gains in the Rickettsiaceae coupled to the widespread RP seen in wolbachiae (**Fig. 6B**). We posit that this model is more parsimonious than RP as an ancestral rickettsial trait, given that RP-inducing genes are mostly found embedded within mobile genetic elements (e.g., WO prophage in *Wolbachia*, RAGE ICEs and plasmids in *Rickettsia* spp.).

RP-inducing genes are found in greater numbers and diversity in wolbachiae, undoubtedly due to their strong association with EAMs of WO prophage genomes and mobile elements (30, 65). In our previous work, we uncovered a link between the characterized CI-inducing CinA/B and CidA/B operons of wolbachiae and a putative TA operon encoded on the pLbAR plasmid of *Rickettsia felis* str. LSU-Lb, an obligate endosymbiont of the booklouse *Liposcelis bostrychophila* (48). We posited that this TA operon induces parthenogenesis in the booklouse: pLbAR is unknown in other strains of *R*. *felis* which do not induce parthenogenesis, in booklice or other arthropods (48), and booklice lacking *R*. *felis* str. LSU-Lb reproduce sexually (128, 129). Aside from being a CndB toxin (harboring both the NUC and DUB domains of wolbachiae CinB and CidB toxins, respectively) the large pLbAR toxin (3,241 aa) was found to carry several other domains present in other *Wolbachia* proteins, most of which are also EAM-associated. These domains, together with the **large central domain** (**LCD**) of the pLbAR toxin, were utilized to identify an extraordinarily diverse array of candidate RP-inducing toxins in a wide-range of intracellular bacteria, some of which are known reproductive parasites (67). Importantly, because this large assemblage of proteins was identified using sequence characteristics of the pLbAR toxin, we consider the collective family linked within the portion of the intracellular mobilome driving the recurrent evolution of RP in invertebrate-borne bacteria.

Aside from CI induction by CinA/B and CidA/B operons (24–27), virtually nothing is known about the functions of related candidate RP-inducing genes. The function of the CinA/B-like operon of *O. tsutsugamushi* is unclear*. O. tsutsugamushi* is the pathogen causing scrub typhus and is capable of inducing parthenogenesis in *Leptotrombidium* mites (130, 131). Given this fact, exploration of its corresponding CinA/B module is warranted. Though the connection to RP in mites is unclear given that CinA/B only occurs in the Boryong strain; a myriad of other proteins encoding the pLbAR LCD are amplified in all *Orientia* genomes. Genome sequences are needed (but not yet available) to determine toxin profiles for *Rickettsia* species either in ancestral lineages (132–134) or Transitional Group rickettsiae (132, 135, 136) that induce parthenogenesis in eulophid wasps or trogiid booklice. Genome sequences are also lacking for *Rickettsia* species from ancestral lineages known to induce male-killing in buprestid and ladybird beetles (137–140). We previously mined an unpublished genome of the male killer *Rickettsia* parasite of the ladybird beetle *Adalia bipunctata* (141) and found a toxin highly similar in architecture to the *Spiroplasma* male killer toxin (142). While both toxins carry an **ovarian tumor** (**OTU**) cysteine protease domain (143), we found a hodgepodge of toxin architectures carrying OTU domains from bacteria known to induce other RP phenotypes, such as parthenogenesis (*w*Fol (69)) and feminization (*w*VulC (144)). Thus, it is clear that additional high-quality (closed) genomes are sorely needed to correlate RP-inducing protein domains and RP phenotypes. New correlations will lead to downstream functional analyses of target’s abilities to induce RP.

A large number of candidate proteins from the pLbAR-derived pool occur in species that are not known as reproductive parasites. These include CndB toxins, NUC- and DUB-domain containing proteins that are much larger than characterized CinB and CidB toxins, diverse OTU-domain containing toxins, and especially proteins harboring the LCD that may or may not exhibit toxin-like profiles (67). Some of these candidate toxins appear to have cognate antidotes, and some genomes carry antidotes only, highlighting the complex diversity of this biological function. Recently, an EAM-encoded transcriptional factor in *w*Mel termed WO-mediated killing (*wmk*) was shown to induce male-specific lethality during early embryogenesis when expressed transgenically in fruit flies (29). This male-killer gene was not detected as part of our pLbAR-derived pool, indicating that RP-inducing gene arsenals are probably much larger than have been described thus far, particularly considering that *cin*, *cid* and *cnd* loci are not present in the genomes of *Cardinium* and *Arsenophonus* reproductive parasites (66, 67, 145) for which the mechanisms for RP induction remain unknown (146, 147).

Candidate RP-inducing toxins with similar features to the pLbAR toxin (large, modular, and associated with mobile genetic elements) are found in several *Wolbachia* genomes (e.g., *w*Fol, *w*Str, *w*DacB, *w*VulC, *w*Con, *w*Ced, *w*CauA). They are often in conjunction with other *cin* and/or *cid* loci, yet not always associated with WO prophage. Consistent with prior reports identifying LGT between *Rickettsia* and *Wolbachia* genomes (21, 48, 148, 149), similar toxins in *Rickettsia* genomes (*R*. *felis* str. LSU-Lb, *R*. *gravesii*, male killer of *Adalia bipunctata*) indicate a dynamic platform for disseminating RP-inducing genes across diverse rickettsial species that includes *Rickettsia* plasmids and WO prophage. Growing numbers of identified *Rickettsia* plasmids (48), proliferative RAGE ICEs in *Rickettsia* and *Orientia* genomes (21, 117, 150–153), and the recent discovery of the first *Wolbachia* plasmid (154), attest to a highly dynamic mobilome not previously realized for obligate intracellular bacteria. The *w*CfeJ fission of a CndB-like toxin into a CinB toxin described in the current study (**Fig. 3D**) supports the premise that larger toxins recombine into WO prophage and undergo streamlining into discrete inducers of CI or other RP phenotypes. As with the evolution of biotin utilization, Rickettsiales provides a framework to study the nature of RP in a diverse array of bacteria that enjoy different relationships with their invertebrate and vertebrate hosts.

## Conclusion

Fleas are understudied vectors of several human and animal diseases (155). While wolbachiae (mostly unnamed strains) have been detected in many flea species (50, 51, 156), it is pressing from a biocontrol perspective to know if these bacteria affect the ability of fleas to vector pathogens. The cat flea transmits pathogenic bacteria such as *Rickettsia typhi* (murine typhus), *R*. *felis* (murine typhus-like illness), *Bartonella* spp. (e.g., cat-scratch disease and various other bartonelloses), and to a lesser extent the plague agent *Yersinia pestis* (157–160). Cat fleas are also intermediate hosts of the tapeworm *Dipylidium caninum* and the filarial nematode *Acanthocheilonema reconditum*, both of which can infect humans (161–163). Other organisms observed in *C*. *felis* include undescribed Baculoviridae, amoebae, trypanosomatids, cephaline gregarines (Apicomplexa), and microsporidia (164). However, nearly all epidemiological studies have focused on determining the frequency and distribution of only certain pathogens (e.g., *Bartonella* spp., *R*. *typhi*, *R*. *felis*), with few documenting wolbachiae co-occurrence with these pathogens (165, 166).

The remarkable opposing features of *w*CfeT and *w*CfeJ offer testable hypotheses for contrasting host relationships (nutritional symbiosis versus RP, respectively). They also provide a framework to study the dynamics of *Wolbachia* co-infections, both in colonies and wild populations. The *w*CfeT and *w*CfeJ genomes yield lessons on the role of LGT in shaping host associations, highlighting the ephemeral nature of nutritional-based mutualism and RP over a general rickettsial platform of obligate intracellular parasitism. Host genome sequences provide clues to on-going mutualism and historical RP, and our hypothesis for host CI-gene capture leading to suppression of RP illuminates possible limitations for deployment of *Wolbachia* reproductive parasites to combat arthropod transmission of human pathogens. In summary, our robust characterization of two divergent strains of *C*. *felis*-associated wolbachiae, as well as the *C. felis* genome itself (45), provides essential information for determining the impact of these bacteria on cat flea biology and pathogen vectoral dynamics.

## Materials and Methods

### Assembly and annotation of *w*CfeT and *w*CfeJ

Our recent report detailed 1) the assembly of *Wolbachia* reads generated from *C*. *felis* genome sequencing, 2) genome circularization and naming of two novel strains (*w*CfeT and *w*CfeJ), and 3) complete genome annotation (45). All relevant information pertaining to *w*CfeT and *w*CfeJ is available at NCBI under Bioproject PRJNA622233.

### Wolbachiae Phylogenomics

Protein sequences (n=84,836) for 66 sequenced *Wolbachia* genomes plus 5 additional Anaplasmataceae (*Neorickettsia helminthoeca* str. Oregon, *Anaplasma centrale* Israel, *A. marginale* Florida, *Ehrlichia chaffeensis* Arkansas, and E. *ruminantium Gardel*) were either downloaded directly from NCBI (n=48), retrieved as genome sequences from the NCBI Assembly database (n=13), contributed via personal communication (n=8; Michael Gerth, Oxford Brookes University), or sequenced here (n=2) (**Table S1**). For genomes lacking functional annotations (n=15), gene models were predicted using the RAST v2.0 server (n=12) or GeneMarkS-2 v1.10_1.07 (n=3; (167)). Ortholog groups (n=3,038) were subsequently constructed using FastOrtho, an in-house version of OrthoMCL (168), using an expect threshold of 0.01, percent identity threshold of 30%, and percent match length threshold of 50% for ortholog inclusion. A subset of single-copy families (n=12) conserved across at least 60 of the 66 genomes were independently aligned with MUSCLE v3.8.31 (169) using default parameters, and regions of poor alignment were masked with trimal v1.4.rev15 (170) using the “automated1” option. All modified alignments were concatenated into a single data set (2,803 positions) for phylogeny estimation using RAxML v8.2.4 (171), under the gamma model of rate heterogeneity and estimation of the proportion of invariant sites. Branch support was assessed with 1,000 pseudo-replications. Final ML optimization likelihood was -52350.085098.

### BOOM characterization

#### Phylogeny estimation

Genomes (n=93) containing at least 4 of the 6 biotin synthesis genes were identified using Blastp searches against the NCBI nr database (e-value < 0.01; accessed January 9, 2020). Protein sequences for these loci were downloaded from NCBI, aligned with MUSCLE v3.8.31 (169) using default parameters, and regions of poor alignment masked with trimal v1.4.rev15 (170) using the “automated1” option. All modified alignments were concatenated into a single data set (1,457 positions) for phylogeny estimation using RAxML v8.2.4 (171), under the gamma model of rate heterogeneity and estimation of the proportion of invariant sites. Branch support was assessed with 1,000 pseudo-replications. Final ML optimization likelihood was -127140.955066. See **Table S2** for NCBI protein accession IDs for all sequences included in this analysis.

#### Gene neighborhood analysis of wCfeT BOOM

Genes in a 25-gene window around the wCfeT BOOM cluster were used to query (Blastp) the NCBI nr database (accessed January 9, 2020). The top 50 matches to each sequence (e-value < 1, BLOSUM45 matrix) were binned by taxonomic assignment as “Wolbachia”, “other Rickettsiales”, or “other (non-Rickettsiales) taxa.” Other (non-Rickettsiales) taxa were further binned by major taxonomic group to assess the boundaries and diversity of the *w*CfeT BOOM MGE.

#### Synteny analysis of biotin genes in wolbachiae

Rickettsiales genomes with at least 5 of the 6 biotin synthesis genes (n=28) were downloaded from NCBI and ortholog groups (n=2,527) constructed with FastOrtho, an in-house version of OrthoMCL (168), using an expect threshold of 0.01, percent identity threshold of 30%, and percent match length threshold of 50% for ortholog inclusion. OGs containing biotin synthesis genes were identified using *Rickettsia buchneri* str. Wikel sequences as seeds. Genome locations for all genes were retrieved from NCBI in gene file format (gff) and used to order the orthologs in each genome.

#### Phylogenetic estimation of the EamA protein of wCfeT

Taxa were selected from Blastp searches against the NCBI Rickettsiales and nr (excluding Rickettsiales) databases using the *w*CfeT EamA protein (WP_168464633.1) as a query. Proteins were aligned with MUSCLE v3.8.31 (169) using default parameters, with regions of poor alignment masked using Gblocks (172). Phylogeny was estimated with the WAG substitution model using RAxML v8.2.4 (171) under the gamma model of rate heterogeneity and estimation of the proportion of invariant sites.

### CI gene characterization

#### Wolbachiae

*w*CfeJ CinA and CinB were compared to other RP-inducing TA operons following our previous approach (67). Another novel CndAB operon was identified on a small scaffold (NCBI Nucleotide ID CABPRJ010001055.1) in the aphid *Cinara cedri*, which has an associated *Wolbachia* symbiont, *w*Ced (173), and was also characterized in a similar manner. Briefly, EMBL’s Simple Modular Architecture Research Tool (SMART) (174) and /or the Protein Homology/analogY Recognition Engine V 2.0 (Phyre2) (175) were used to predict the following domains: OTU cysteine protease (Pfam OTU, PF02338) (143), endoMS-like (176) and NucS-like (177) PD-(D/E)XK nuclease (Pfam PDDEXK_1, PF12705) (178), CE clan protease (Pfam Peptidase_C1, PF00112) (179), Latrotoxin-CTD (180), and ankyrin repeats. Individual protein schemas were generated using Illustrator of Biological Sequences (181) with manual adjustment.

#### C. felis

A *cidA*-like gene was identified in the *Ctenocephalides felis* genome during initial genome decontamination (45). Briefly, a contig composed of a mosaic of flea- and *Wolbachia*-like sequence was identified by our comparative BLAST-based pipeline and flagged for manual inspection. Gene models on this contig were predicted using the RAST v2.0 server. The resulting proteins were used to create a custom Blast database that was queried using Blastp with CinAB sequences from *Wolbachia* endosymbiont of *Culex quinquefasciatus* Pel (CAQ54390, CAQ54391), and CidAB sequences *Wolbachia pipientis* wAlbB (CCE77512, CCE77513). The contig was subsequently confirmed as belonging to the largest scaffold (NW_020538040) in the *C. felis* assembly. A second, *cinA*-like gene was identified by querying all CDS in the published *C. felis* assembly (NCBI accession ASM342690v1) with the same set of *Wolbachia* CinAB and CidAB sequences.

### Screening colonies and wild populations of cat fleas for *w*CfeT and *w*CfeJ

Upon receipt from collaborators, cat fleas were stored at −20°C in EtOH until processing for DNA extraction and qPCR. Fleas were sexed under a stereomicroscope and surfaced sterilized for 5 min with 1% bleach solution, followed by 5 min with 70% EtOH, and finally with three washes using molecular grade water. Individual fleas were flash-frozen in liquid N_2_ and ground to powder with sterile pestle in a 1.5ml microcentrifuge tube. DNA was extracted from dissected tissues using the Zymo Quick-DNA Miniprep plus kit following the solid tissue protocol (Zymo Research; Irvine, CA). The presence of *w*CfeT, *w*CfeJ, and *C. felis* DNA was determined via qPCR using the following primer sets: Cfe#18S|179, 5’-TGCTCACCGTTTGACTTGG-3’ and 5’-GTTTCTCAGGCTCCCTCTCC-3’ (182); *w*CfeT#ApaG|75: 5’-GCCGTCACTGGCAGGTAATA-3’ and 5’-GCTGTTCTCCAATAACGCCA-3’, *w*CfeJ#CinA|76, 5’-AGCAACACCAACATGCGATT-3’ and 5’-GAACCCCAGAGTTGGAAGGG-3’. Each primer set amplicon was sequenced before use to ensure specificity. qPCR was performed using a QuantStudio 3 with the PowerUp SYBR green mastermix in a 20μl reaction volume containing 2μl of sample DNA with 400nM of each primer. Cycling parameters were consistent with the “fast cycling mode” including a 2 min cycle at 50°C, a 2 min cycle at 95°C, 40 cycles with 3 sec at 95°C and 30 seconds at 60°C followed by a melt curve analysis. Standard curves were generated to quantify the number of DNA copies present in each reaction. Results were expressed as the ratio of *Wolbachia* amplicons to *C. felis* 18S rDNA copies to normalize for varying efficiency of DNA extractions. Fleas were considered positive for wolbachiae if the flea 18S rDNA Ct value is 30 or lower and the *Wolbachia* Ct value is 37 or lower.

### Determining *w*CfeT and *w*CfeJ localization in *C*. *felis* tissues

Elward Laboratories (EL; Soquel, CA) cat fleas stored at −80°C were used to assay tissue localization of *w*CfeT and *w*CfeJ. Ovaries and midguts of individual female fleas were dissected under a stereomicroscope. DNA extraction and qPCR were performed as described above.

### Data availability

All relevant information pertaining to *w*CfeT and *w*CfeJ is available at NCBI under Bioproject PRJNA622233. Complete genome sequences are available in the NCBI RefSeq database at accession identifiers NZ_CP051156.1 (*w*CfeT) and NZ_CP051157.1 (*w*CfeJ).

## SUPPLEMENTAL MATERIAL

Data as well as custom analysis scripts generated for this project that are not published elsewhere are provided on GitHub in the “*w*Cfe_genomes” repository available at https://www.github.com/wvuvectors/wCfe_genomes.

**TABLE S1.** Information supporting the *Wolbachia* genome-based phylogeny estimation.

**TABLE S2.** Information supporting the phylogeny estimation of biotin genes.

**TABLE S3.** Information supporting the comparative analysis of BOOM gene neighborhoods in wolbachiae genomes.

**FIG S1.** Phylogeny estimations of biotin synthesis enzymes and EamA transporters.

(A) Complete phylogeny estimation of BOOM and other *bio* gene sets from diverse bacteria. Tree was estimated from the concatenation of six *bio* enzymes (BioC, BioH, BioF, BioA, BioD, and BioB) or subsets in certain cases (see **Table S2** for all sequence information and **Materials and Methods** for details on dataset processing and tree estimation). Branch support was assessed with 1,000 pseudo-replications. Final ML optimization likelihood was -127140.955066. In taxon color scheme, [R] denotes Rickettsiales, with Holosporaceae [H] as a revised family of Rhodospirillales (101). The families of Rickettsiales are similarly noted: Anaplasmataceae [A], Midichloriaceae [M], Rickettsiaceae [R], and “*Candidatus* Deianiaeaceae” [D], with the latter considered provisional (91).

(B) Estimated phylogeny of EamA transporters. See **Materials and Methods** for details on dataset processing and tree estimation. Branch support was assessed with 1,000 pseudo-replications. Final ML optimization likelihood was -23058.797227. Taxon color scheme as described in panel **A**.

**FIG S2.** *In silico* characterization of the *w*CfeT prophage genome.

Schema shows the comparison of the *w*CfeT and WOVitA1 prophage genomes, with gene descriptions following prior studies (26, 30, 148, 183). *w*CfeT genes within the typical Eukaryotic Association Module (EAM) were used as queries in Blastp searches against the NCBI nr protein database, with summary statistics provided for top subjects (dashed box).

**FIG S3.** Synteny analysis of biotin synthesis genes in Rickettsiales genomes.

Rickettsiales taxa with at least 5 of 6 biotin synthesis genes (n=28) were analyzed for co-linearity of their biotin genes. Ortholog groups (n=2,527) were constructed from these genomes as described for Fig. 1 (see **Materials and Methods**) and OGs containing biotin synthesis genes identified. Genome locations were retrieved from NCBI in gene file format (gff) and used to order the orthologs in each genome. For unclosed genomes, all contigs were included in OG construction; only a single genome (*Candidatus* Aquarickettsia rohweri) contained biotin synthesis genes on multiple contigs. Biotin genes are colored according to the ideal BOOM shown at the top. Black boxes indicate stretches of unrelated genes. Taxa are grouped into 3 classes according to the synteny of biotin genes in their genomes: Class I taxa demonstrate completely conserved gene order; Class II taxa contain only a single gene out of order (Class IIA: BioA; Class IIB: BioB); Class III taxa exhibit little or no conservation of gene order. All blocks are anchored on BioB for ease of comparison, except Class IIB which are anchored on BioF instead.

**FIG S4.** Bioinformatics analysis of BioU and related methyl esterases.

(A) Alignment of *Ehrlichia chaffeensis* and *Anaplasma centrale* BioU proteins with orthologs from other Rickettsiales species. BioU orthologs were retrieved from Blastp searches against the NCBI Rickettsiales databse using *E*. *chaffeensis* str. Sapulpa BioU (EAM85518.1) as a query. The catalytic residues (Ser–Asp–His) characteristic of methyl esterases are highlighted yellow (184). Sequences were aligned with MUSCLE v3.8.31 (169) using default parameters. NCBI protein accession numbers are listed in panel **C**.

(B) Percent identity matrix calculated from the alignment shown in panel **A**.

(C) Estimated phylogeny of proteins shown in panel **A**. Two alphaproteobacterial outgroups and two hymenopteran mitochondrial methyl esterases are included. Phylogeny was estimated using RAxML v8.2.4 (171) under the gamma model of rate heterogeneity and estimation of the proportion of invariant sites. Branch support was assessed with 1,000 pseudo-replications. Final ML optimization likelihood was - 5448.767799.

(D) Analysis of eight rickettsial genomes that contain biotin synthesis genes except for the BioH methyl esterase. Blastp searches against the rickettsial genomes were conducted using the following queries, all of which have been shown to orthogonally substitute for BioH: BioV of *Helicobacter* sp. L8b (TSA87024.1); BioG of *Haemophilus influenzae* str. CGSHiCZ412602 (AIB45959.1); BtsA of *Moraxella catarrhalis* str. CCUG 353 (KZR94829.1); BioJ of *Francisella marina* str. E103-15 (QEO56965.1); BioZ of *Mesorhizobium japonicum* str. R7A (AAG47795.1); EstN1 of *Nitrososphaera viennensis* str. EN76 (AIC14436.1); BioK of *Prochlorococcus marinus* str. SS51 (KGG31655.1).

## ACKNOWLEDGMENTS

We acknowledge Dr. Michael Gerth (University of Liverpool) for sharing unpublished contigs from sequencing projects for *Wolbachia* endosymbiont of *Zootermopsis nevadensis* (*w*Zoo), *Wolbachia* endosymbiont of *Mengenilla moldrzyki* (*w*Men), and *Wolbachia* endosymbiont of *Ctenocephalides felis* (*w*Cte). We are grateful to Dr. Michael Dryden (Kansas State University), Dr. Bill Donahue (Sierra Research Laboratories, Modesto, CA), Dr. Lance Durden (Georgia Southern University), and Eric Dotseth (Division of Infectious Disease Epidemiology, West Virginia Department of Health and Human Resources) for contributing cat fleas.

This work was supported with funds from the National Institutes of Health National Institute of Allergy and Infectious Diseases grants (R21AI26108 and R21AI146773 to JJG, R01AI017828 and R01AI126853 to AFA, and R01AI122672 to KRM). TPD and VIV were supported by start-up funding provided to TPD by West Virginia University. JFB was supported by start-up funds from Auburn University, USDA Hatch Grant (1015922) and an Alabama Agricultural Experiment Station SEED grant. The content is solely the responsibility of the authors and does not necessarily represent the official views of the funding agencies. The funders had no role in study design, data collection and analysis, decision to publish, or preparation of the manuscript.

