## Supplemental Figures for "Evolution of *Wolbachia* Mutualism and Reproductive Parasitism: Insight from Two Novel Strains that Co-infect Cat Fleas"

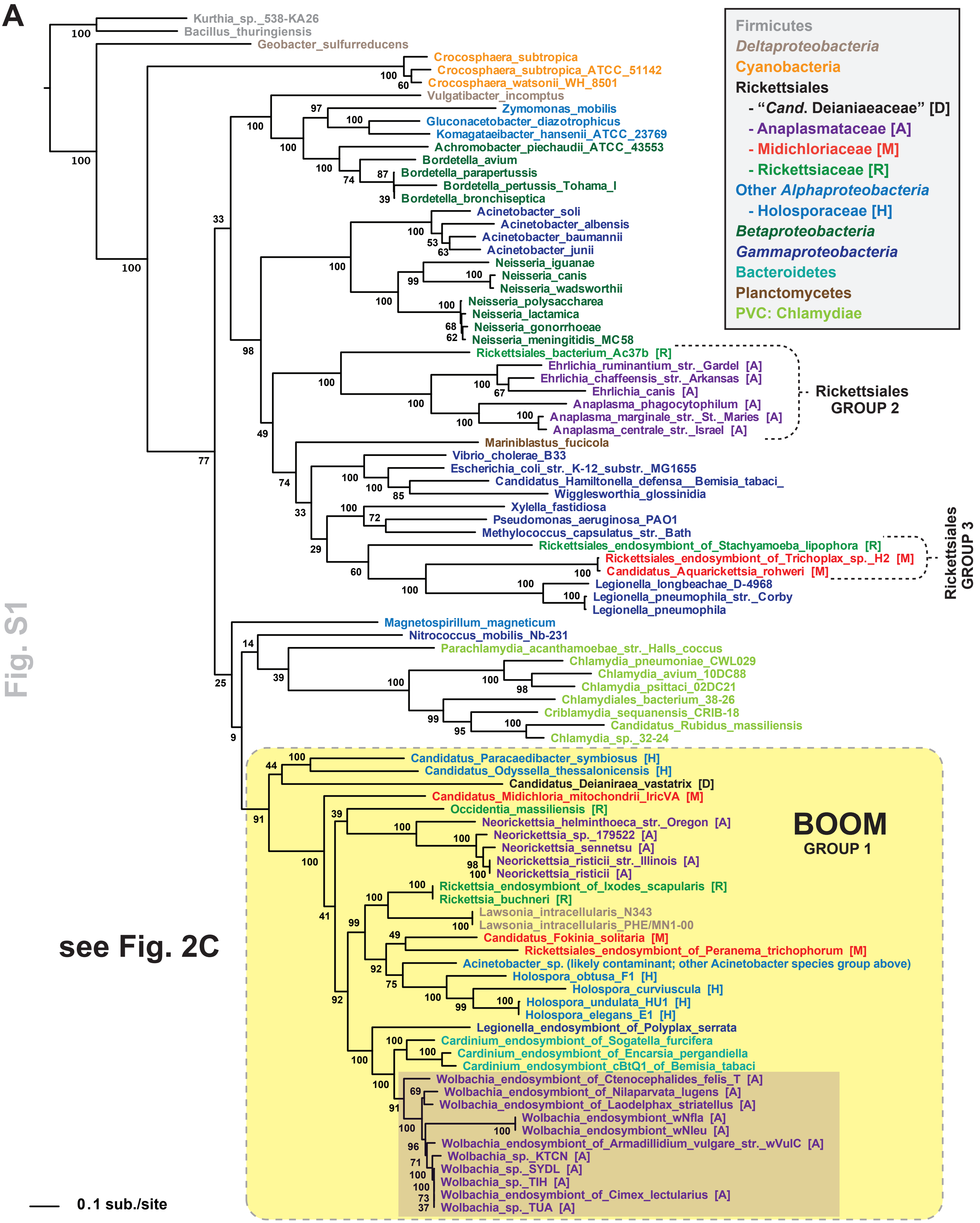


see Fig. 2C

Fig. S1

**B**

**Fig. S1**

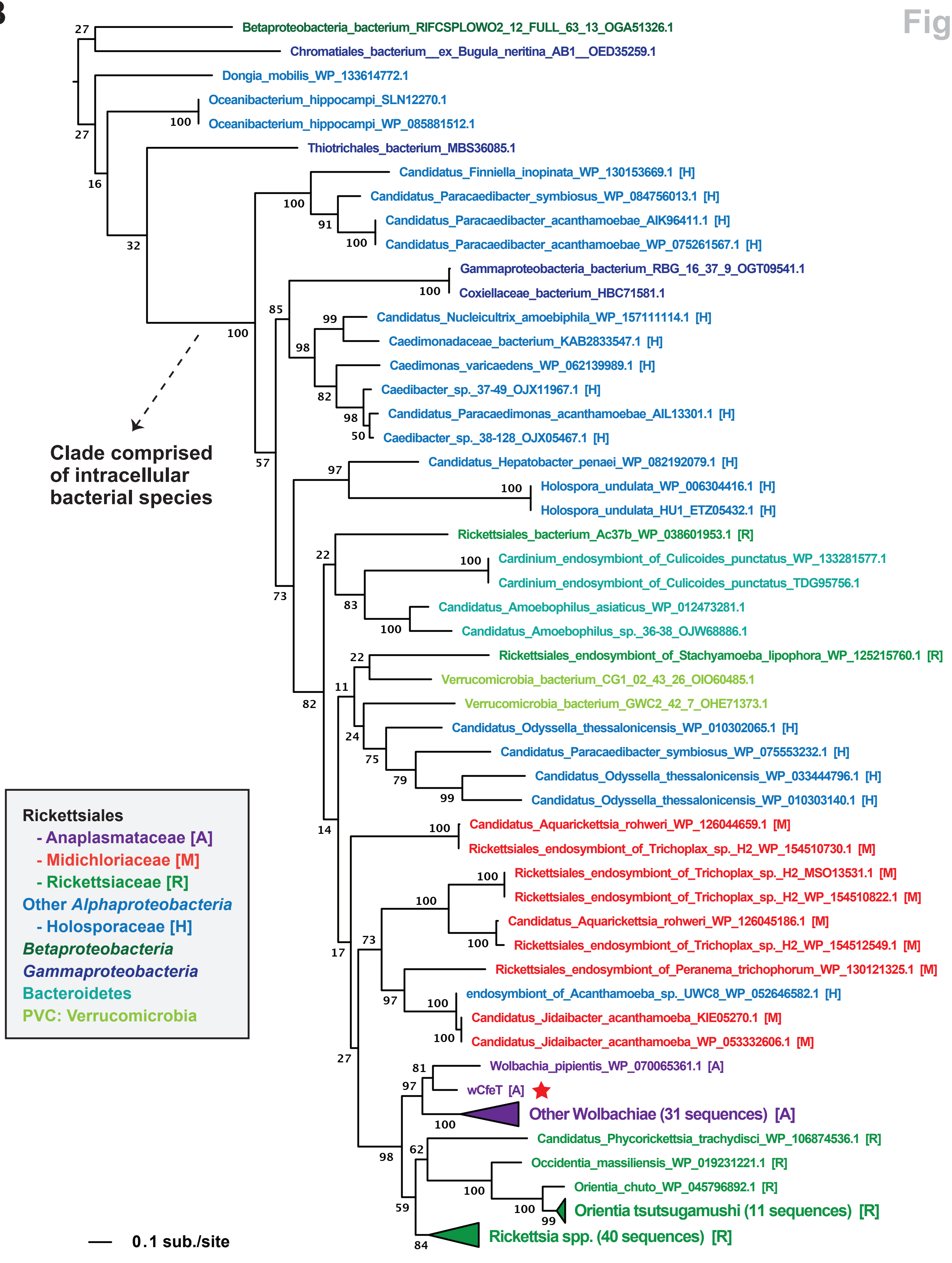
Betaproteobacteria_bacterium_RIFCSPLOWO2_12_FULL_63_13_OGA51326.1
Chromatiales_bacterium__ex_Bugula_neritina_AB1__OED35259.1
Dongia_mobilis_WP_133614772.1
Oceanibacterium_hippocampi_SLN12270.1
Oceanibacterium_hippocampi_WP_085881512.1
Thiotrichales_bacterium_MBS36085.1
Candidatus_Finniella_inopinata_WP_130153669.1 [H]
Candidatus_Paracaedibacter_symbiosus_WP_084756013.1 [H]
Candidatus_Paracaedibacter_acanthamoebae_AIK96411.1 [H]
Candidatus_Paracaedibacter_acanthamoebae_WP_075261567.1 [H]
Gammaproteobacteria_bacterium_RBG_16_37_9_OGT09541.1
Coxiellaceae_bacterium_HBC71581.1
Candidatus_Nucleicultrix_amoebiphila_WP_157111114.1 [H]
Caedimonadaceae_bacterium_KAB2833547.1 [H]
Caedimonas_varicaedens_WP_062139989.1 [H]
Caedibacter_sp._37-49_OJX11967.1 [H]
Candidatus_Paracaedimonas_acanthamoebae_AIL13301.1 [H]
Caedibacter_sp._38-128_OJX05467.1 [H]
Clade comprised of intracellular bacterial species
Candidatus_Hepatobacter_penaei_WP_082192079.1 [H]
Holospora_undulata_WP_006304416.1 [H]
Holospora_undulata_HU1_ETZ05432.1 [H]
Rickettsiales_bacterium_Ac37b_WP_038601953.1 [R]
Cardinium_endosymbiont_of_Culicoides_punctatus_WP_133281577.1
Cardinium_endosymbiont_of_Culicoides_punctatus_TDG95756.1
Candidatus_Amoebophilus_asiaticus_WP_012473281.1
Candidatus_Amoebophilus_sp._36-38_OJW68886.1
Rickettsiales_endosymbiont_of_Stachyamoeba_lipophora_WP_125215760.1 [R]
Verrucomicrobia_bacterium_CG1_02_43_26_OIO60485.1
Verrucomicrobia_bacterium_GWC2_42_7_OHE71373.1
Candidatus_Odyssella_thessalonicensis_WP_010302065.1 [H]
Candidatus_Paracaedibacter_symbiosus_WP_075553232.1 [H]
Candidatus_Odyssella_thessalonicensis_WP_033444796.1 [H]
Candidatus_Odyssella_thessalonicensis_WP_010303140.1 [H]
Candidatus_Aquarickettsia_rohweri_WP_126044659.1 [M]
Rickettsiales_endosymbiont_of_Trichoplax_sp._H2_WP_154510730.1 [M]
Rickettsiales_endosymbiont_of_Trichoplax_sp._H2_MSO13531.1 [M]
Rickettsiales_endosymbiont_of_Trichoplax_sp._H2_WP_154510822.1 [M]
Candidatus_Aquarickettsia_rohweri_WP_126045186.1 [M]
Rickettsiales_endosymbiont_of_Trichoplax_sp._H2_WP_154512549.1 [M]
Rickettsiales_endosymbiont_of_Peranema_trichophorum_WP_130121325.1 [M]
endosymbiont_of_Acanthamoeba_sp._UWC8_WP_052646582.1 [H]
Candidatus_Jidaibacter_acanthamoeba_KIE05270.1 [M]
Candidatus_Jidaibacter_acanthamoeba_WP_053332606.1 [M]
Wolbachia_pipientis_WP_070065361.1 [A]
wCfeT [A]
Other Wolbachiae (31 sequences) [A]
Candidatus_Phycorickettsia_trachydisci_WP_106874536.1 [R]
Occidentia_massiliensis_WP_019231221.1 [R]
Orientia_chuto_WP_045796892.1 [R]
Orientia tsutsugamushi (11 sequences) [R]
Rickettsia spp. (40 sequences) [R]
Rickettsiales
- Anaplasmataceae [A]
- Midichloriaceae [M]
- Rickettsiaceae [R]
Other Alphaproteobacteria
- Holosporaceae [H]
Betaproteobacteria
Gammaproteobacteria
Bacteroidetes
PVC: Verrucomicrobia
0.1 sub./site

| *w*CfeT proteins | | Top Blastp hit | | | | | | | |
|---|---|---|---|---|---|---|---|---|---|
| Protein | Size | Accession | Size | Annotation | Taxon | Max, tot. | Cov. | E value | %ID |
| WP_168464803 | 309 | APR98706 | 311 | Patatin-like phospholipase | *w*Fol | 514, 514 | 100% | 0.0 | 78.14% |
| WP_168464804 | 72 | AZU37351 | 74 | ParD-like family antidote | *w*Bta | 107, 107 | 97% | 7e-29 | 72.86% |
| WP_168464805 | 96 | OAM00647 | 92 | RelE/ParE family toxin | *w*DacA | 154, 154 | 95% | 5e-47 | 80.43% |
| ---------------------- | 44 | ---------------- | ---- | ------------------------------------------------------------- | ---------- | --------------- | -------- | -------- | ----------- |
| WP_168464806 | 262 | APR98707 | 257 | PHA03095 (Ank repeats) | *w*Fol | 321, 321 | 98% | 1e-107 | 60.31% |
| WP_168464807 | 82 | APR99034 | 86 | XRE family transcriptional regulator | *w*Fol | 111, 111 | 97% | 5e-30 | 68.75% |
| WP_168464808 | 648 | APR98618 | 651 | DNA ligase (NAD(+)) LigA | *w*Fol | 1007, 1007 | 99% | 0.0 | 73.92% |
| WP_168464809 | 79 | APR97852 | 96 | XRE family transcriptional regulator | *w*Fol | 139, 139 | 98% | 4e-41 | 89.74% |
| WP_168464414 | 545 | WP_143688845 | 493 | Group II intron RevTranscriptase/maturase | *w*Str | 992, 992 | 89% | 0.0 | 99.59% |
| WP_168464810 | 310 | APR98606 | 312 | Recombination-promoting nuclease/put. tnp | *w*Fol | 506, 506 | 100% | 3e-179 | 79.35% |
| WP_168464811 | 301 | APR98615 | 302 | Hypothetical protein (put. rhoptry protein) | *w*Fol | 421, 421 | 100% | 9e-146 | 74.17% |
| WP_168464812 | 226 | APR98945 | 217 | DNA repair protein RadC | *w*Fol | 367, 367 | 96% | 7e-127 | 84.86% |

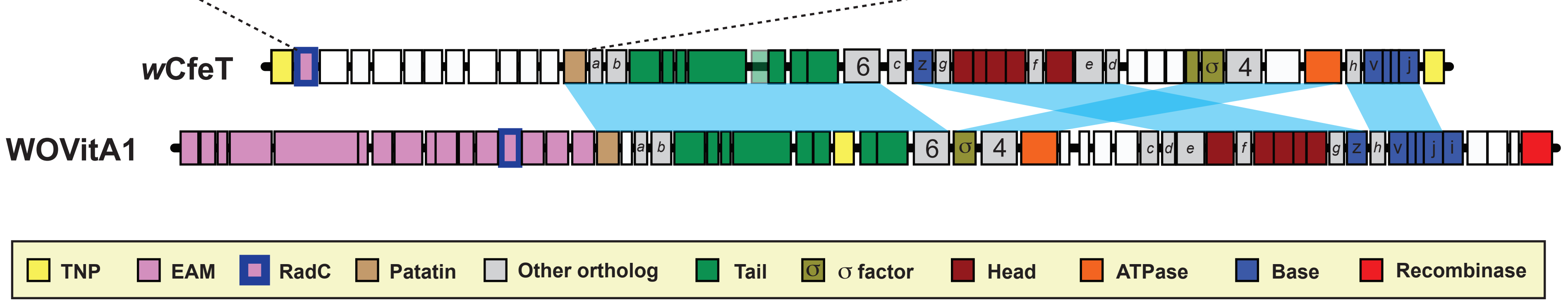


Fig. S2

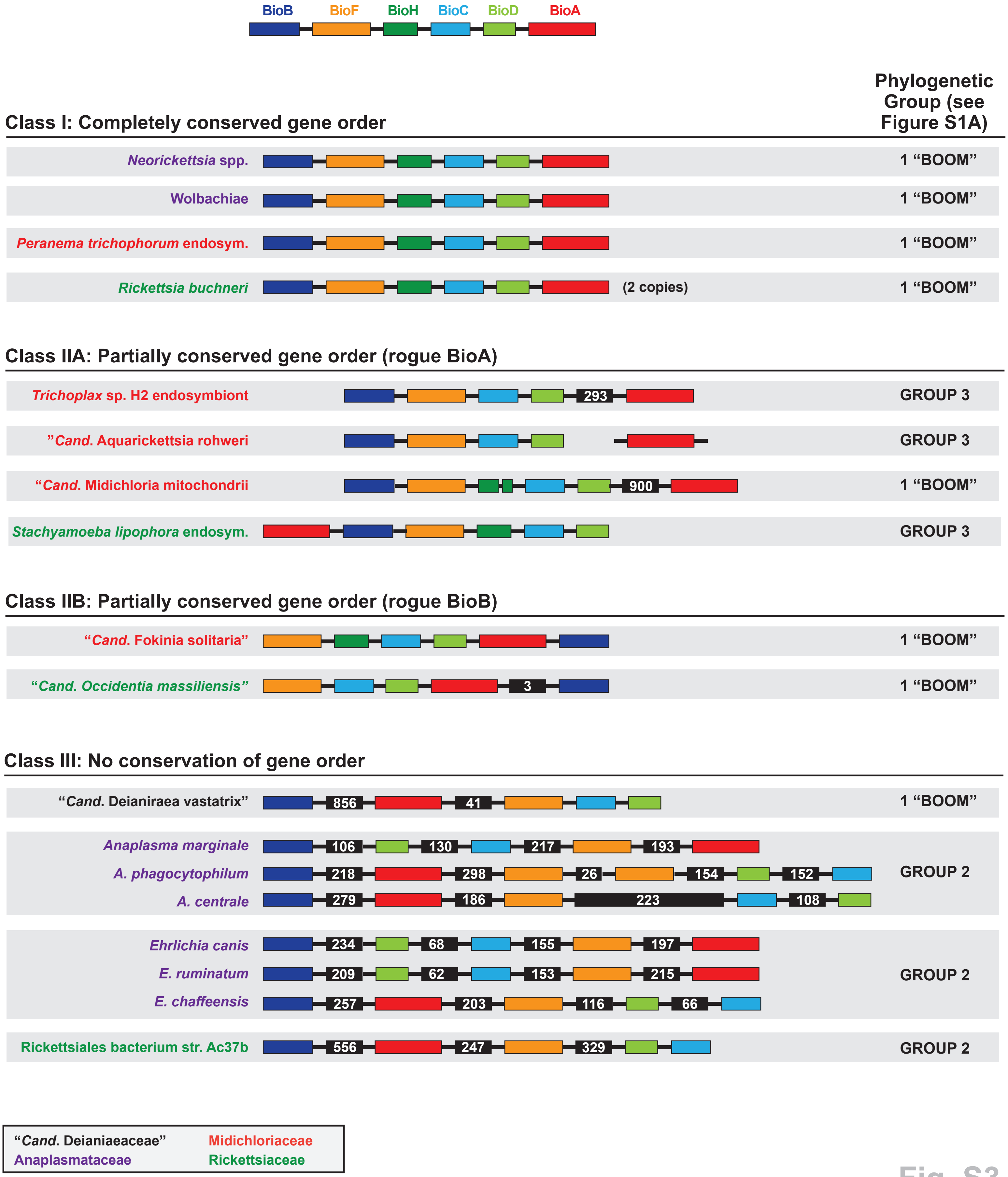

BioB
BioF
BioH
BioC
BioD
BioA
Phylogenetic Group (see Figure S1A)
Class I: Completely conserved gene order
Neorickettsia spp.
1 “BOOM”
Wolbachiae
1 “BOOM”
Peranema trichophorum endosym.
1 “BOOM”
Rickettsia buchneri
(2 copies)
1 “BOOM”
Class IIA: Partially conserved gene order (rogue BioA)
Trichoplax sp. H2 endosymbiont
293
GROUP 3
”Cand. Aquarickettsia rohweri
GROUP 3
“Cand. Midichloria mitochondrii
900
1 “BOOM”
Stachyamoeba lipophora endosym.
GROUP 3
Class IIB: Partially conserved gene order (rogue BioB)
“Cand. Fokinia solitaria”
1 “BOOM”
“Cand. Occidentia massiliensis”
3
1 “BOOM”
Class III: No conservation of gene order
“Cand. Deianiraea vastatrix”
856
41
1 “BOOM”
Anaplasma marginale
106
130
217
193
A. phagocytophilum
218
298
26
154
152
A. centrale
279
186
223
108
GROUP 2
Ehrlichia canis
234
68
155
197
E. ruminatum
209
62
153
215
E. chaffeensis
257
203
116
66
GROUP 2
Rickettsiales bacterium str. Ac37b
556
247
329
GROUP 2
“Cand. Deianiaeaceae”
Midichloriaceae
Anaplasmataceae
Rickettsiaceae


Fig. S3

A

```
U ┤ Ehrlichia chaffeensis              ------MSDHYK--LVLSNPQGLHITYRQLLGN----KASIIFFGGFNSNMQGTKATALY    48
  ┤ Anaplasma centrale                 MLRQSCVGAVVGEEKRLELGNSSYISYMQTTVQS---PVSVVFFGGFMSDMHGTKAQHLF    57
Rickettsiales bacterium Ac37b          ------MNIAFN-VQKLLLSNNNYIAYSKINSKTQNELPGIIFFSGFNSNMQGTKARNLT    53
Neorickettsia helminthoeca             ------MQTKYL------DTSHGRIAYQTFDNNP---EVGVLFMTGLASDMSGRKSERLR    45
Rickettsia buchneri                    ------MHKLYN------KTQDKFIVYDNYRIINTN-IPSVIFLHGLMSSMKSTKAIYLI    47
wCle                                   -------MDYCK----LFGESGKYIAYRKLQGR----RTSIVFFGGFASNMNGTKATAIY    45
● wPip_Pel                             -------MNYCK----LLDGSNGHIAYRKLQGK----KASIVFFSGFASNMDGTKATAVY    45
● Neoehrlichia lotoris                 ------MTDYSPSFLVLDHNNNHKIAYRQLKKNSN--LPSILLLGGFGSNMYGEKATALY    52
● Rickettsia typhi                     ------MHKLYN------KTQDKFIVYDNYRIINTN-IPSVIFLHGLMSSMQSTKAIYLI    47
● Orientia tsutsugamushi Boryong       ------MHKIYT------LDDGRFIAYRQHKSQKNS-LINIIFLHGMMSNMSGKKSSYLY    47
                                                                  .    * *            .:::: *: *.* . *:  :

                                                                                          Ser
U ┤ Ehrlichia chaffeensis              DYCKSHNLGLILFDYLGHGQSDGQFTDYNISDWYKNCIEIITQLTPTNRPKIIIGSSMGA    108
  ┤ Anaplasma centrale                 EYCKSHGVHCTVFDYFGHGSSSGEFQECTISDWYASCVSVVESL--TSAPLVIVGSSMGG    115
Rickettsiales bacterium Ac37b          EYCQNNNYNFIKFDYLGHGLSSGIFHECTIGIWLENCLSIIDNL--TTDKHIFIGSSMGG    111
Neorickettsia helminthoeca             SFCENNQVAFTRFDYFGHGRSEGSFLHGNISKWTENALEVLERV--TTGKQILVGSSMSG    103
Rickettsia buchneri                    DYCKKNNYNFIVFDNFGHGNASGQFEDQTISDWLEGVSLILDKL--IDKEAILVGSSMGG    105
wCle                                   KFCQENDVALVLFDYFGHGHSSGDFTDYTISDWQKNCTRVINEL--TSSKQIIIGSSMGG    103
● wPip_Pel                             KFCQENDIALVLFDYFGHGNSSGDFADYTISDWQKNCAKVISEL--TSNKQIIIGSSMGG    103
● Neoehrlichia lotoris                 NYCNKHNLNLTVFDYLGHGHSSGNFTDYTIGDWYKNCISVIESL--TNGPQIIIGSSMGG    110
● Rickettsia typhi                     DYCKKNNYNFIVFDNFGHGNAYGQFEDQTISDWLEGVALILDKL--IETEAILIGSSMGG    105
● Orientia tsutsugamushi Boryong       QLCQEEDLNFLAFDNYGHGNSSGRFIDQTIESWFDATRAIMYHTS-NNFKNIIVGSSLGG    106
                                       . *:.         **   *** : * *    .*   *        ::           :::***:..

U ┤ Ehrlichia chaffeensis              WLMLLVAISHQDKVSHLISLAGAPDFTESLIFQKLNTQQKDELYKYGQITL--SQNSNNM    166
  ┤ Anaplasma centrale                 WLMLLTALSHGRRVRGLVGMAPAPDFTESL---DLSESQRAEMMRTGKTIK--NTDNC--    168
Rickettsiales bacterium Ac37b          WLALLASILRPEKVAGIICIAAAPDFTENLIWNTLSLEEKNKLQTQGIIKL--SSNYCEG    169
Neorickettsia helminthoeca             WMMFKIAEKHPEKVKGLVGIAAAPDFTED-FLEGLTHETKQALEKNGYFTF--TRNRDE-    159
Rickettsia buchneri                    WLALLAALRFPDKIKGLVCVAPAPDFTEN-IWQNISLNDQNKMQKEGILEV--SGKNCEH    162
wCle                                   WLMLLTALQIPERIAALIGVSSAPDFTEDLIFKQLSGKQKEELDSKGVVDF--TSGRC--    159
● wPip_Pel                             WLMLLTALQFPEKIAALIGISSAPDFTEDLIFKQLSGKQKEELGSKGVIDF--TSEHC--    159
● Neoehrlichia lotoris                 WLMLLIAQSYPHKVISLLGLAPAPDFTENLIFNKLTQEQKDCLHTNNQIIF--TFNKYED    168
● Rickettsia typhi                     WLALLAALRFPDKIKCLICVAPALDFTEN-IWQNISLNDQNKMKKEGIIEV--SSENCQH    162
● Orientia tsutsugamushi Boryong       WLAMLAAIKNEIEISGVVALAPAIDFTETLIWNKLTEKNKNSMIHTGYIELGGTGNTCNN    166
                                       *: :  :       :  :: :: * ****       :. . .  :    .        .

                                                                              Asp
U ┤ Ehrlichia chaffeensis              YSYVITRNLIEDGRKHLLLNQESINITCPITLIHGMNDDTVPYQTSITVAEKIKSDNVNL    226
  ┤ Anaplasma centrale                 -SYVITKKLIDDGKAHLLMNKREIAVECPMVLIHGMDDTVVPYQVSLEIAGKVKSGDVRV    227
Rickettsiales bacterium Ac37b          -EYEISLKLIEEAREHLILNK-PLDIKCPIYLLHGMADKDVPYNFSLDLVNSISSQDITV    227
Neorickettsia helminthoeca             -KLVITKTLLDDGKKNLILTQ-RIKVPCPVVLLHGLADDIVSYRKSIELAELIESSPVEV    217
Rickettsia buchneri                    -KYPISYKLIEDAKKHLLLTKKQIDINTPLHIIHGMLDEDVPYNVSVKLLEKITSKQIVM    221
wCle                                   -AYKITKNLIEDGRKKLLLNKETIDINSPVRLLHSINDKDVPYQTSLNLAERVKSTDVEV    218
● wPip_Pel                             -AYKITKNLIEDGRKNLLLNREAIDINCPVRLLHSINDKDVPYQTSLNLAEKIKSTDVEV    218
● Neoehrlichia lotoris                 RSYDITDNLIKDGRKHLLLNNDNININCPVILIHSMSDLVVPYSTSIHVAEKITSTNVNL    228
● Rickettsia typhi                     -KYPISYKLIEDAKKHLLLRKQQIEINIPVHIIHGMLDKNVPYNVSVKLLEKITSKQIVM    221
● Orientia tsutsugamushi Boryong       -KYHISYNLICNARKYLLLNKPTINIQCPIAIIHGMQDQEVPYQGSIDLINKVQTHYSTL    225
                                           *: .*: :..  *:: .   : :   *: ::*.: *  *.*   *: :    : :    :

                                         His
U ┤ Ehrlichia chaffeensis              HLIKSANHNLSDDTSLNIIFKYIKEAVEQSIQVK    260
  ┤ Anaplasma centrale                 HLSKSGTHRLTDEHSLGLMLESVKGLMRPSVPV-    260
Rickettsiales bacterium Ac37b          KLVKDAGHGMSTKVNLYLLYNTINELITKITNK-    260
Neorickettsia helminthoeca             RLIKGADHSMSDPTSITVLTDTVRALI-------    244
Rickettsia buchneri                    KLIKDGHHNLSREEDLKVMTNSLEEVISLSNIK-    254
wCle                                   HLTKSAEHNMSDSHSLKILFQAIREFLPTV----    248
● wPip_Pel                             HLIKSAEHNMSDNHSLKILFKTIREFLPGEIYN-    251
● Neoehrlichia lotoris                 HLIKSGNHYLRDEHSLNVTFSAIQSLLAQC----    258
● Rickettsia typhi                     KLIKDGNHNLSRKEDLKVIANSLEEMISNIK---    252
● Orientia tsutsugamushi Boryong       KLLKYADHFLSDSVSLSHISYAIKEIINARLV--    257
                                       .* *  . * :     .:        :   :
```

**U** = characterized BioU

(shaded box) = taxa with BioH

● = no biotin synthesis

Fig. S4

Fig. S4

**B**

| | Otsu | Nhel | Acen | Rtyp | Rbuc | R37b | wCle | wPip_Pel | Nlot | Echa |
|---|---|---|---|---|---|---|---|---|---|---|
| *Orientia tsutsugamushi* Boryong (Otsu) | 100.00 | | | | | | | | | |
| *Neorickettsia helminthoeca* (Nhel) | 31.56 | 100.00 | | | | | | | | |
| U *Anaplasma centrale* (Acen) | 34.01 | 37.08 | 100.00 | | | | | | | |
| *Rickettsia typhi* (Rtyp) | 40.48 | 31.97 | 35.92 | 100.00 | | | | | | |
| *Rickettsia buchneri* (Rbuc) | 41.50 | 34.43 | 38.87 | 88.49 | 100.00 | | | | | |
| Rickettsiales bacterium Ac37b (R37b) | 41.11 | 37.30 | 36.90 | 46.22 | 45.85 | 100.00 | | | | |
| *w*Cle | 38.21 | 40.25 | 44.90 | 37.14 | 36.73 | 44.13 | 100.00 | | | |
| *w*Pip_Pel | 37.10 | 43.15 | 43.15 | 39.43 | 37.50 | 46.80 | 84.27 | 100.00 | | |
| *Neoehrlichia lotoris* (Nlot) | 40.24 | 38.93 | 45.82 | 37.60 | 37.60 | 41.57 | 52.02 | 52.82 | 100.00 | |
| U *Ehrlichia chaffeensis* (Echa) | 41.67 | 37.45 | 45.02 | 41.37 | 39.44 | 41.96 | 52.82 | 54.98 | 60.24 | 100.00 |

**C**

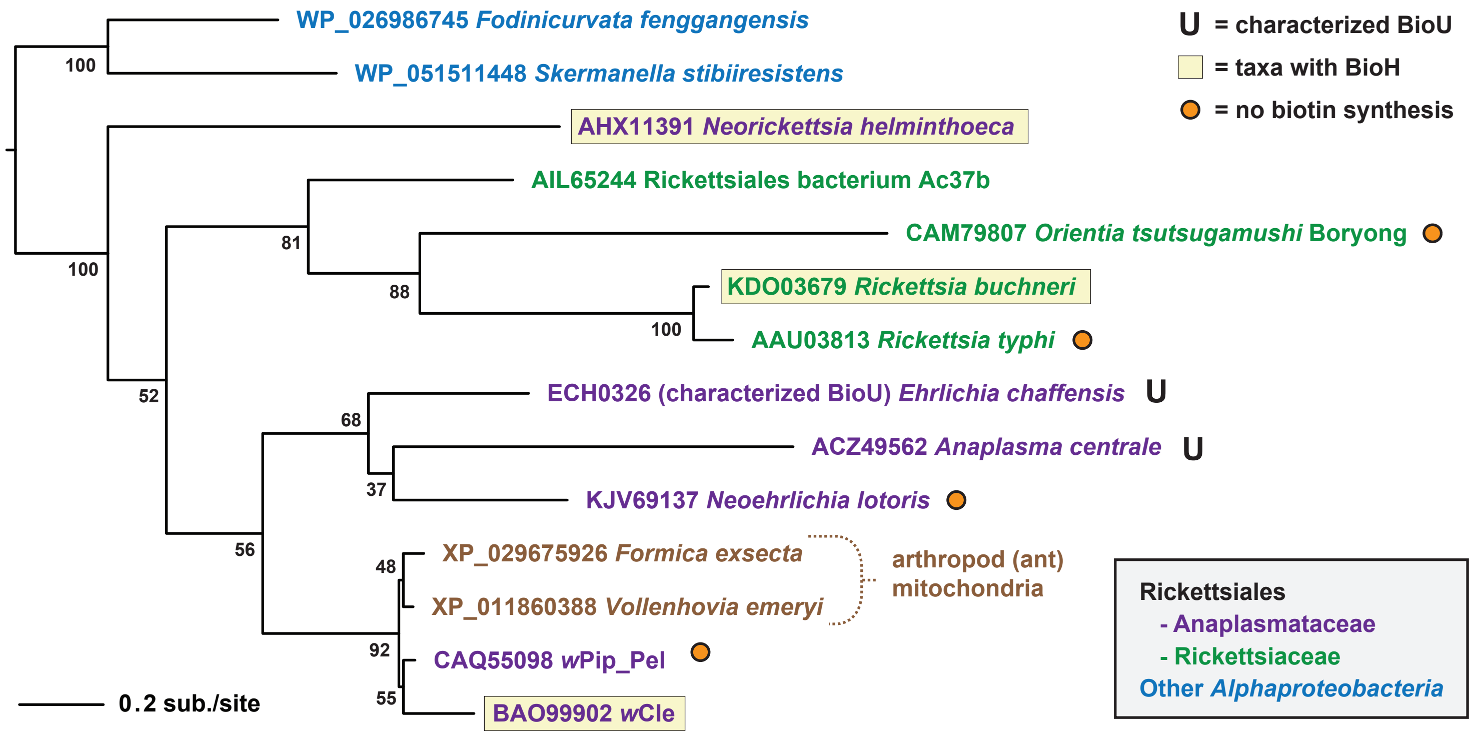


**D**

| NCBI taxid | Taxon | Blastp results (%ID) BioV | BioG | BtsA | BioJ | BioZ * | EstN1 | BioK | BioU |
|---|---|---|---|---|---|---|---|---|---|
| 768 | Anaplasma | NSS | NSS | NSS | NSS | 33 | ~27; (2) | NSS | Present |
| 943 | Ehrlichia | NSS | NSS | NSS | NSS | 33 | ~27; (2) | NSS | Present |
| 2021221 | Rickettsiales endosymbiont of Trichoplax sp. H2 | NSS | NSS | NSS | NSS | 33 | NSS | NSS | NSS |
| 2602574 | Ca. Aquarickettsia rohweri | NSS | NSS | NSS | NSS | 33 | NSS | NSS | NSS |
| 1528098 | Rickettsiales bacterium Ac37b | NSS | NSS | NSS | NSS | 35 | 31; AIL65750 | NSS | Present |
| 752179 | Occidentia massiliensis | NSS | NSS | NSS | NSS | 35 | NSS | NSS | NSS |
| 2163644 | Ca. Deianiraea vastatrix | NSS | NSS | 37; (1) | NSS | 30 | NSS | NSS | NSS |

(1) QED23496 (5-methyltetrahydropteroyltriglutamate--homocysteine S-methyltransferase); only 12% query coverage

* % similarity to FabH; BioZ is similar to FabH but was able to complement *E. coli ΔbioH* mutants (PMID: 11320134).

(2) similarity to a cohort of MhpC-like Abhydrolase 5 proteins (unrelated to BioU; *E. chaffeensis* Arkansas protein is ECH_0221).
